# Saving large fish through harvest slots outperforms the classical minimum-length limit when the aim is to achieve multiple harvest and catch-related fisheries objectives

**DOI:** 10.1101/738708

**Authors:** Robert N. M. Ahrens, Micheal S. Allen, Carl Walters, Robert Arlinghaus

## Abstract

We address the problem of optimal size-selective exploitation in an age-structured fish population model by systematically examining how density- and size-dependency in growth, mortality and fecundity affect optimal harvesting patterns when judged against a set of common fisheries objectives. Five key insights are derived. First, while minimum-length limits often maximize the biomass yield, exploitation using harvest slots (i.e., regulations that protect both immature and very large individuals) can generate within 95% of maximum yield, and harvest slots generally maximize the number of fish that are harvested. Second, density-dependence in growth and size-dependent mortality predict more liberal optimal size-limits than those derived under assumptions of no density and size-dependence. Third, strong density-dependence in growth maximizes the production of trophy fish in the catch only when some modest harvest is introduced; the same holds for numbers harvested, when the stock-recruitment function follows the Ricker-type. Fourth, the inclusion of size-dependent maternal effects on fecundity or egg viability has only limited effects on optimal size limits, unless the increase in fecundity with mass (“hyperallometry”) is very large. Extreme hyperallometry in fecundity also shifts the optimal size-limit for biomass yield from the traditional minimum-length limit to a harvest slot. Fifth, harvest slots generally provide the best compromises among multiple objectives. We conclude that harvest slots, or more generally dome-shaped selectivity to harvest, can outperform the standard minimum-length selectivity. The exact configuration of optimal size limits crucially depends on objectives, local fishing pressure, the stock-recruitment function and the density and size-dependency of growth, mortality and fecundity.

## Introduction

Traditional fish population and harvesting theory has largely been developed under a single management objective - to maximize biomass yield (Beverton & Holt, 1957; Schaefer, 1957), translated into the long-term goal of directing fishing mortality to levels that guarantee the maximum sustainable yield (MSY) (Larkin, 1977). Single species age-structured population models widely support the prediction that biomass yield is maximized by implementing sharp selectivity in harvest and direct exploitation on ages/sizes where a cohort’s biomass peaks (Froese et al., 2016). One common harvest regulation able to safeguard such selectivity is a minimum-length limit (Allen, Hanson, Ahrens, & Arlinghaus, 2013; Clark, Alexander, & Growing, 1980; Jensen, 1981; Maceina, Bettoli, Finely, & DiCenzo, 1998; Ricker, 1945) or more generally size- and/or age-dependent sigmoidal selectivity where small and young fish are protected from harvest and large and old(er) fish are harvested (Beverton & Holt, 1957; van Gemert & Andersen, 2018). Management of size-selectivity enjoys substantial support among stakeholders and is particularly prevalent in fisheries where direct management and control of fishing mortality rate, e.g. through quotas on landings or effort controls, is impossible, too costly, or logistically daunting. Examples are recreational fisheries (Arlinghaus, Lorenzen, Johnson, Cooke, & Cowx, 2016; Noble & Jones, 1999) and data-limited fisheries in the developing world (Prince & Hordyk, 2019; Wolff, Taylor, & Tesfaye, 2015).

Depending on fishing mortality rates, implementation of minimum-length limits may come at the cost of severe age and size truncation, leading to a strong decline or even loss of large and by the same token old fish in exploited stocks (Arlinghaus, Matsumura, & Dieckmann, 2010; Barnett, Branch, Ranasinghe, & Essington, 2017; Beamish, McFarlane, & Benson, 2006; Pierce, 2010). The demise of large fish may negatively affect fishing quality, in particular in fisheries where the catch or harvest of large fish produces particularly large benefits to humans (Asche, Chen, & Smit, 2015; Beardmore, Hunt, Haider, Dorow, & Arlinghaus, 2015; Carlson, 2016; Witteveen, in press). There are also fisheries where intermediate fish sizes (“plate-size fish” or “kitchen fish”) generate higher market prices than either smaller or larger individuals (Reddy et al., 2013), suggesting that harvest slots - combinations of minimum and maximum size limits or generally dome-shaped selectivity to harvest - may be superior harvesting patterns than minimum-length limits under certain fisheries objectives and conditions (Arlinghaus et al., 2010; Gwinn et al., 2015; Law, 2007; Ayllón et al. 2019).

Indeed, depending on local culture and stakeholder composition (e.g., the mixture of commercial and recreational fishers in a local fishery) biomass yield maximization may not be socially optimal (Johnston, Arlinghaus, & Dieckmann, 2010; Johnston, Beardmore, & Arlinghaus, 2015). In particular recreational anglers often value other fisheries objectives more strongly than biomass yield, e.g., the catch of memorable large fish or high catch rates, from which only a portion is taking home for dinner (Arlinghaus et al., 2019). For harvest-oriented recreational fisheries, Gwinn et al. (2015) argued that the number of fish harvested, rather than biomass yield, maybe a more suitable target as higher numbers of acceptably sized fish available for distribution among a large pool of anglers may produce higher overall utility than a maximized biomass yield where the landings are composed by on average larger, but overall much fewer fish (Arlinghaus et al., 2010; Ayllón et al., 2019). Many local fisheries are co-exploited by both commercial and recreational fisheries, e.g., most coastal fisheries (Arlinghaus et al., 2019). Here, both biomass oriented (tailored towards commercial fishers) and more catch or size oriented fisheries objectives (tailored towards recreational fisheries) will be jointly important. A key question then is which size limit to use to produce acceptable compromises that fulfill a range of often-conflicting objectives without perhaps optimizing any single one (Ayllón et al., 2019; García-Asorey, Escati-Penaloza, Parma, & Pascual, 2011; Gwinn et al., 2015; Koehn & Todd, 2012). Gwinn et al. (2015) presented a single-species age-structured model that suggested harvest slots could constitute such a compromise regulation that maybe superior to classical minimum-length limits in meeting several objectives jointly.

Minimum-length limits have recently come under scrutiny because of conservation concerns associated with strong juvenescence effects (e.g., Anderson et al, 2008; Arlinghaus et al., 2010; Birkeland & Dayton, 2005; Sánchez-Hernández, Shaw, Cobo, & Allen, 2016). Strong declines in highly fecund, large and old fish under intensive fishing may reduce total egg output (Barneche, Robertson, White, & Marshall, 2018; Berkeley, Hixon, Larson, & Love, 2004a; Froese 2004; Hsieh, Yamauchi, Nakazawa, & Wang, 2010) and has been reported empirically and in models to destabilize stock dynamics (Anderson et al., 2008; Botsford, Holland, Field, & Hastings, 2014; Hixon, Johnson, & Sogard, 2014; Hsieh et al. 2006, 2010). Recent work has emphasized that traditional assumptions about the scaling of fecundity with fish size maybe wrong, challenging optimal harvesting patterns derived from traditional harvesting theory (Barneche et al., 2018; Hixon et al., 2014). While a linear increase in fecundity with the mass of individual fishes (so called isometric size-fecundity relationship) is well established in fisheries ecology and a standard assumption in stock assessments (Walters & Martell, 2004), recent studies have suggested two types of often-unaccounted size-dependent maternal effects to be widespread in fish. First, hyperallometric relationships of fish mass and fecundity across a vast range of fish species result in a positive relationship of mass-specific fecundity (i.e., relative fecundity in eggs per female mass) and body weight (Barneche et al., 2018). Barneche et al. (2018) in turn proposed that not accounting for hyperallometry in fecundity might foster inappropriate fisheries policies that focus harvest on particularly valuable old and large, highly fecund fish. Secondly, some fish species have been shown to have elevated egg and larval qualities with increasing size and by the same token age and body mass (e.g., Arlinghaus et al., 2010; Berkeley, Chapman, & Sogard, 2004b; Bravington, Grewe, & Davies, 2016; Hixon et al., 2014; Venturelli et al., 2010). However, the relevance of both the hyperallometric fecundity reserve associated with large sizes as well as size-dependent maternal effects on offspring quality for population dynamics and optimal harvesting are matters of substantial debate (Andersen, Jacobsen, & van Denderen, 2019; Arlinghaus et al., 2010; Arnold et al., 2018; Berkeley et al., 2004a; Cooper, Barbiéri, Murphy, & Lowerre-Barbieri, 2013; Hixon et al., 2014; O’ Farrell & Botsford, 2006; Shaw, Sass, & VandeHey, 2018). Indeed, as long as spawning stock biomass remains sufficiently high, biological sustainability of the exploited population should be safeguarded under minimum-length limit regulations. One key condition is to set the minimum harvest size above the size at maturation (Myers & Mertz, 1998; Froese, 2004; Prince & Hordyk, 2019) and to control discard mortality (Coggins, Catalano, Allen, Pine, & Walters, 2007; Johnston et al., 2015).

Other conservation and fisheries problems may occur when minimum length limits are set without properly accounting for key aspects of density-dependence and natural mortality rate in fish population regulation (Brousseau & Armstrong, 1987). Published decision-trees about which type of size limit to choose to meet fisheries objectives carefully account for the strength of density-dependent growth and degree of natural mortality (Arlinghaus et al., 2016; FAO, 2012). Yet, most age-structured models that have been used to derive insights into optimal size limit assume no density-dependence in growth and no size-dependency in mortality. For example, the classical yield-per-recruit model of Beverton & Holt (1957) neglects density feedback on individual growth and assumes constant adult mortality. Variants of this model have been intensively studied and used to examine the likely outcomes of a range of minimum-length and other size-based harvest limits in exploited stocks, targeted by both commercial and recreational fisheries (e.g., Campos & Freitas, 2014; Maceina et al., 1998; Sánchez-Hernández et al., 2016; Wolff et al., 2015).

Yet, density-dependence is a key regulating factor of most fish stocks (Rose, Cowan, Winemiller, Myers, & Hilborn, 2001). While most fisheries scientists assume, rightly perhaps (Zimmermann, Ricard, & Heino, 2018), that most density dependence happens through juvenile mortality and recruitment early in life, there is increasing evidence that late-in-life density-dependence in growth (i.e., growth plasticity) can be important in selected stocks (Lorenzen & Enberg, 2002; Lorenzen, 2005; Zimmermann et al., 2018). When density-dependence in growth is strong, particularly in the juvenile stage, minimum-length limits may contribute to “stock piling” and stunting under the length limit (Tesch, 1959), thereby eroding the productivity of exploited stocks (Arlinghaus et al., 2016). Such conditions have been implicated to contribute to the growth depression of young cod *(Gadus morhua}* in the Eastern Baltic (Svendäng & Hornburg, 2014) and to strongly affect the dynamics of exploited freshwater top predators (Andersen, Jacobsen, Jansen, & Beyer, 2017; Gilbert & Sass, 2016; Persson et al., 2003; Tesch, 1959). Density-dependent growth may also limit the production of trophy fish when resource limitation at high biomass density constrains individuals from reaching their full growth potential (Gilbert & Sass, 2016). Under such conditions, some modest harvesting might release the necessary resources to foster growth and achieve attainment of large body sizes. Despite the prevalence of density-dependent growth in fish stocks, few models examine the costs and benefits of length limits explicitly accounting for growth plasticity (Lorenzen, 2016; Zimmermann et al., 2018).

Many single-stock age-structured models also assume constant natural mortality rates in adults (Beverton & Holt, 1957; Froese et al., 2008; Gwinn et al., 2015; Maceina et al., 1998; Sánchez-Hernández et al., 2016). However, size-dependent natural morality is widespread across a wide range of fishes (Andersen, 2019; Lorenzen, 2000; Peterson & Wroblewski, 1984). Together with density-dependent growth, size-dependent mortality can have strong impacts on how fish stocks respond to size-selective harvesting and other fisheries management interventions (Andersen, 2019; Lorenzen, 2005). In particular, strong inverse size-dependent mortality benefits large size over small size, therefore assuming the natural mortality rate *M* is constant across the adult life stage may underestimate the population dynamical consequences that protection of large individuals can have in exploited stocks. Additionally, natural mortality rate scales directly with fish productivity (Garcia, Sparre, & Csirke, 1989), hence stocks with strong size-dependent mortality can likely support larger harvest without collapsing.

Size-limits are unlikely to produce optimal outcomes on all dimensions because there are fundamental trade-offs to navigate as the fish population changes in response to harvest. For example, while biomass yield tends to be maximized at intermediate equilibrium biomass, catch rates are maximal under unexploited conditions when abundance is maximal (Beverton & Holt, 1957). Gwinn et al. (2015) used an age-structured model calibrated to several life-history prototypes ranging from short-lived to long-lived species, suggesting that while minimum length limits maximized biomass yield, harvest slots produced better compromises among the numbers that were harvested and the catch of large, trophy fish. The limitation of the model by Gwinn et al. (2015) relates to the omission of density-dependent growth and size-dependent mortality, and they did not examine the systematic impact of Ricker-type stock recruitment compared to the standard Beverton-Holt stock recruitment model. Other models have accounted for some of these processes, but these model were calibrated to specific species (e.g., northern pike, *Esox lucius,* Arlinghaus et al., 2010; brown trout, *Salmo trutta,* Ayllón et al., 2019; catfishes of the genus *Ictalurus,* Steward, Long, & Shoup, 2016; lake trout *Salvelinus namaycush,* Lenker, Weidel, Jensen, & Solomon, 2006; or walleye, *Sander vitreus,* Moreau & Matthias, 2018). Importantly, none of these studies have systematically asked which harvest policy is optimal given a spectrum of fishery objectives. Instead, discrete size limit configurations were usually modelled in light of single objectives. No study known to the authors has systematically examined the relative performance of sigmoidal and dome-shaped selectivity on fisheries performance across a broad range of fisheries objectives when density-dependent growth, size-dependent mortality and size-dependent hyperallometry in fecundity and egg viability are present to various degrees, and stock recruitment functions vary from Beverton-Holt-type to Ricker-type with cannibalistic feedback. Most of the globe’s fish stocks analysed so far are regulated assuming a Beverton-Holt-type stock-recruitment, but cannibalistic top predators - common targets particularly of anglers - tend to follow the Ricker model (Szuwalski, Vert-Pre, Punkt, Branch, & Hilborn, 2015). Optimal harvesting is likely to be driven by the stock-recruitment relationship, because cannibalistic feedback can constrain the production of offspring in the Ricker model (Ricker, 1954), but not in the Beverton-Holt (1957) model.

The objective of this paper was to systematically examine the impact of density-dependent recruitment and growth as well as size-dependency in mortality and fecundity/egg viability on optimal size-limits across different management objectives. We asked what type of size-selectivity (i.e., which harvest regulation) is optimal when judged against individual objectives (e.g., biomass yield, number of fish harvested or catch rate) and when judged against an integrative multi-objective function designed to achieve compromises across multiple objectives. The former was done to provide evidence for how to possibly manage a system for one type of predominant fisheries stakeholder. The latter approach simulated cases where managers are tasked to jointly suit different stakeholders in one fishery (e.g., commercial and recreational fisheries). Following Gwinn et al. (2015) we hypothesized that harvest slots would produce the best compromise regulation across a wide range of assumptions about density- and size-dependence in growth, mortality and fecundity/viability. We further expected that increasing degree of density-dependent growth and size-dependent mortality would render optimal harvest regulations more liberal and that the presence of hyperallometry in fecundity and size-dependency in egg viability would promote harvest slots to be particularly suited relative to minimum-length limits (Arlinghaus et al., 2010). By evaluating the effects of several life history characteristics that occur in most fish stocks (e.g., size dependent and density dependent growth and mortality, form of the stock recruit curve, etc.), our analysis has broad applicability to management strategies across both recreational and commercial fisheries.

## Material and Methods

### Model species

The prototypical species that we modelled represented the life history of northern pike (hereafter pike). This species was chosen because it represents an aquatic top predator that is widely distributed in Eurasia and North America in both freshwater and low salinity brackish waters (Skov & Nilsson, 2018). The pike and its close relative, the muskellunge *(Esox masquinogy),* constitute prime targets of fisheries throughout their distributional range (Crane et al., 2015). Pike have also colonized brackish coastal ecosystems, e.g., in the Baltic Sea, where they are co-exploited by commercial and recreational fisheries. Co-exploitation means that possibly conflicting fisheries objectives exist for the same stock that appropriate management regulations must compromise. For example, a jointly exploited stock may be desired to be managed both for as high biomass yield as possible to suit commercial fisheries as well as high catch rate to suit the desires of selected recreational anglers.

The northern pike has a few additional features in its life-history that renders it a suitable model to explore for its reaction to size limits. Importantly, pike populations are governed by both intracohort as well as intercohort cannibalism (Persson et al., 2006). Therefore, pike mortality is size-related in both males and females (Haugen et al., 2007). Although cannibalism constitute a key mechanism that should lead to a Ricker-like stock-recruitment relationship (Ricker, 1954), there is uncertainty about which stock-recruitment relationship best describes pike. Published work failed to detect a relationship of spawner stock size and recruitment (Paxton, Winfield, Fletcher, George, & Hewitt, 2009) or reported Ricker-type (Edeline et al., 2008; Minns, Randall, Moore, & Cairns, 1996; Langangen et al., 2011). Pike stocks have been assumed to exhibit density-dependent growth (Arlinghaus, Matsumura, & Dieckmann, 2009), yet other work failed to find evidence for growth plasticity under natural gradients in density (Lorenzen & Enberg, 2002; Mann, 1980). Density-dependence in growth, should it exist, will not only affect the compensatory reserve of pike populations and their resiliency to harvest (Beverton & Holt, 1957), but will also affect mortality through its impact on growth rate and thus size at age (Lorenzen, 2005).

Pike have been reported to show isometric (i.e., linear) increases in fecundity with mass (Frost & Kipling, 1967), but more recent work suggests that gonad mass as well as egg numbers may scale hyperallometrically with mass (Edeline et al., 2007). In addition, there is evidence for size-dependent maternal effects on egg and larval viability in pike, although few studies exist on this topic (Arlinghaus et al., 2010; Kotakorpi et al., 2013). Therefore, there remains substantial uncertainty about the presence of hyperallometry (i.e., increases in relative fecundity with mass) as well as size or mass-dependent maternal effects on egg quality and/or viability in pike, similar to many other species.

### Model formulation

#### General modelling approach

Optimal size regulations over a range of fishery objectives for our pike-like target species were explored using a classical age structured population model (Walters & Martell, 2004) (Figure 1), but with the additional features of including density-dependent population processes affecting both recruitment and growth as well as size dependence in mortality, maturation and egg production. We varied the stock recruitment function from Beverton-Holt to Ricker-type to represent a large family of cases representing the majority of exploited stocks on the globe. We also allowed for the option of fecundity (egg numbers) to scale hyperallometrically with mass as well as the possibility of a positive linear effect of maternal age on egg-to-recruit survival (i.e., egg/larval viability effect). We contrasted predictions of hyperallometry in fecundity with the standard isometric relationships. The fisheries-management objective could be set out of a suite of single objectives (biomass harvested, number harvested, number caught, and trophy sized caught) as well as the combined utility of all single objectives expressed as a linear combination of a logarithmic form of each objective. This log-scale combined utility function ensured that no single objective of the combined utility function was ignored in searching for an optimal compromise solution (Walters & Martell, 2004). Given a specified management objective, either single-objective or combined objective, lower and upper size limits were allowed to vary to maximize the objective given a specified potential fishing intensity (Figure 1), expressed as the instantaneous fishing mortality rate, *F,* in relation to the adult natural mortality, *F/M,* where *M* is the instantaneous natural mortality rate. Note the fishing mortality input into the model was the potential fishing mortality that could be exerted if individuals were fully selected for by the fishery. The ultimate fishing mortality was affected by the selectivity ogive that resulted from the upper and lower size limits selected to maximize a given objective. Given the dependency of the system state on the fishing mortality, including potential discard mortality resulting from the lower and upper size limits as well as the cascading effects resulting from density dependent growth and size-dependent mortality, equilibrium solutions were found by iterative numerical methods. Model equations as well as parameters descriptions and values are presented in Tables 1 and 2 and are further summarized below.

**Figure 1.**
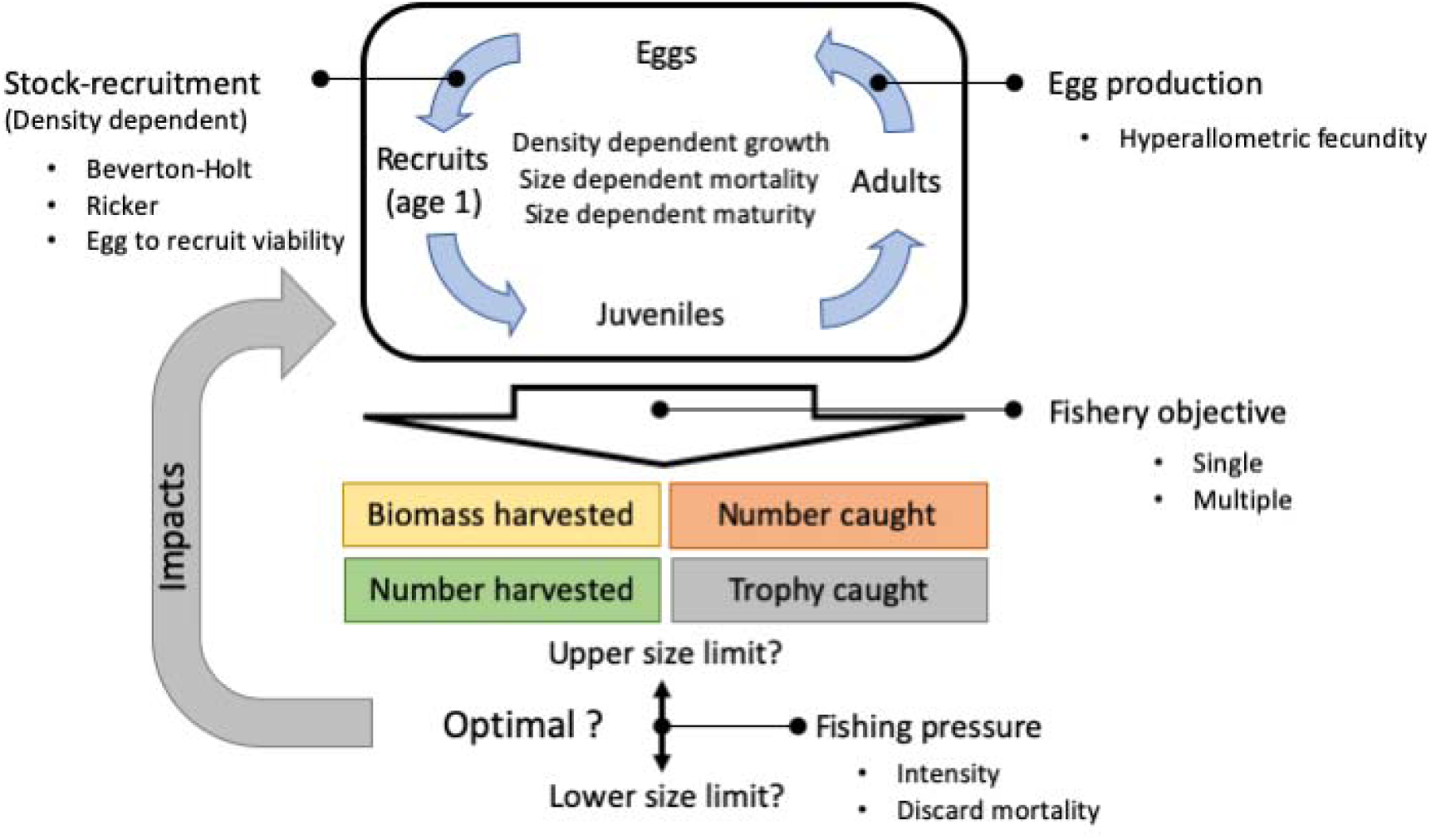
Conceptual description of the age structured fish population model. Bullet points indicate varying processes in the model. The colored boxes indicate management objectives. The question marks indicate the search for the optimal harvest regulations (in terms of size-selectivity) while accounting for complex population dynamical feedbacks resulting from harvesting (“impacts”) until reaching equilibrium.

**TABLE 1.**
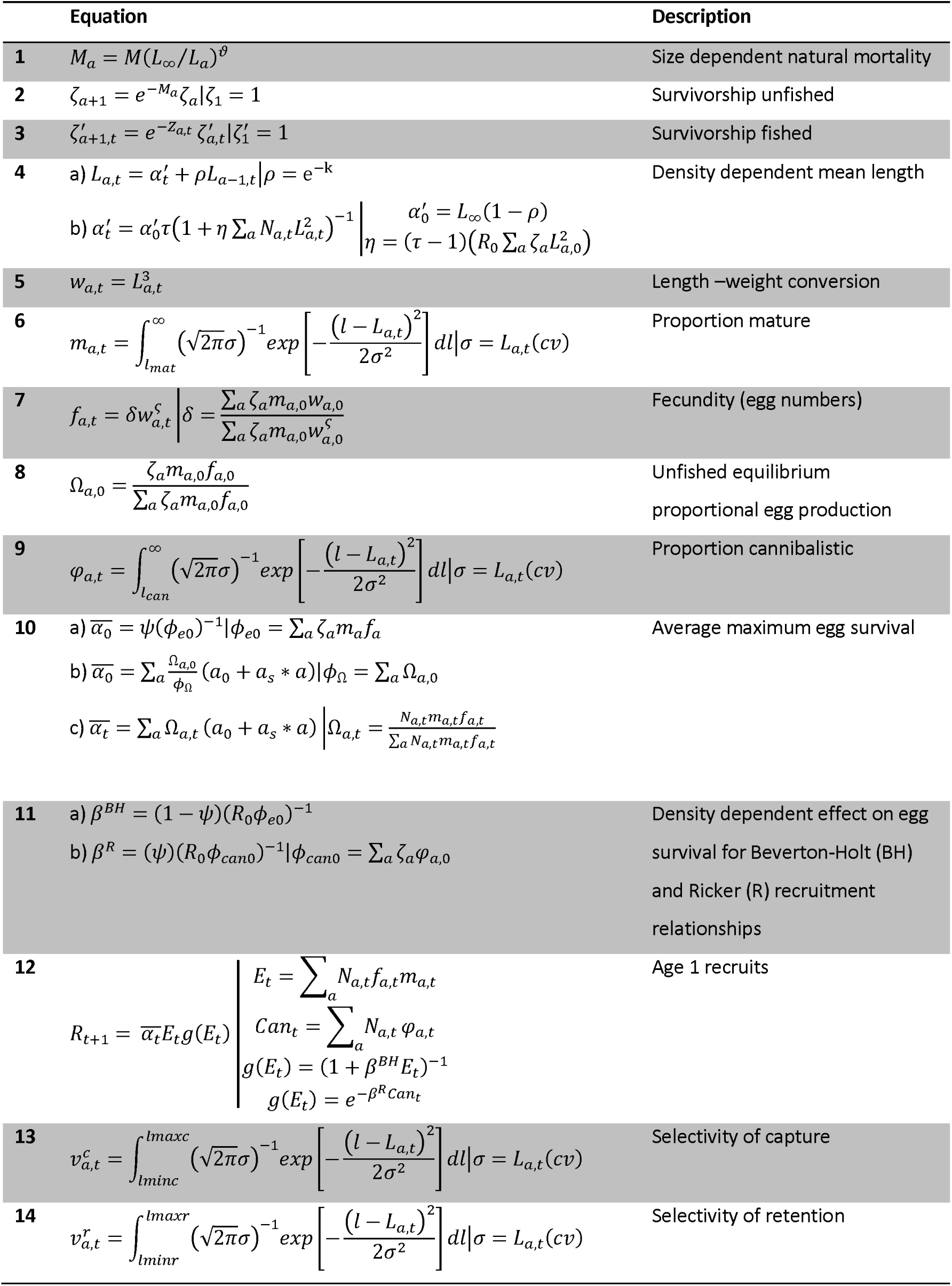
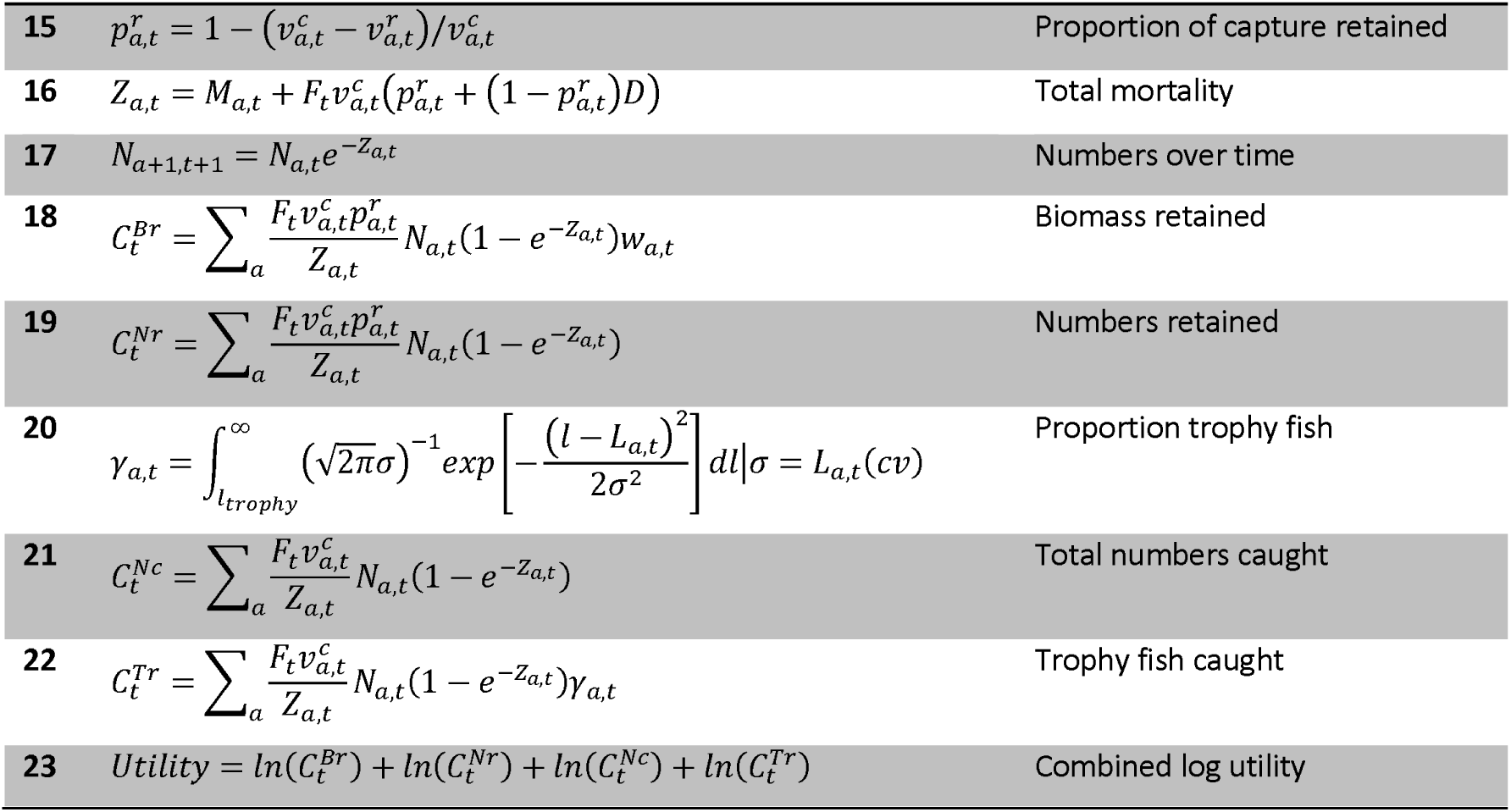
Model equations and description. For description of symbols, see Table 2

**Table 2.**
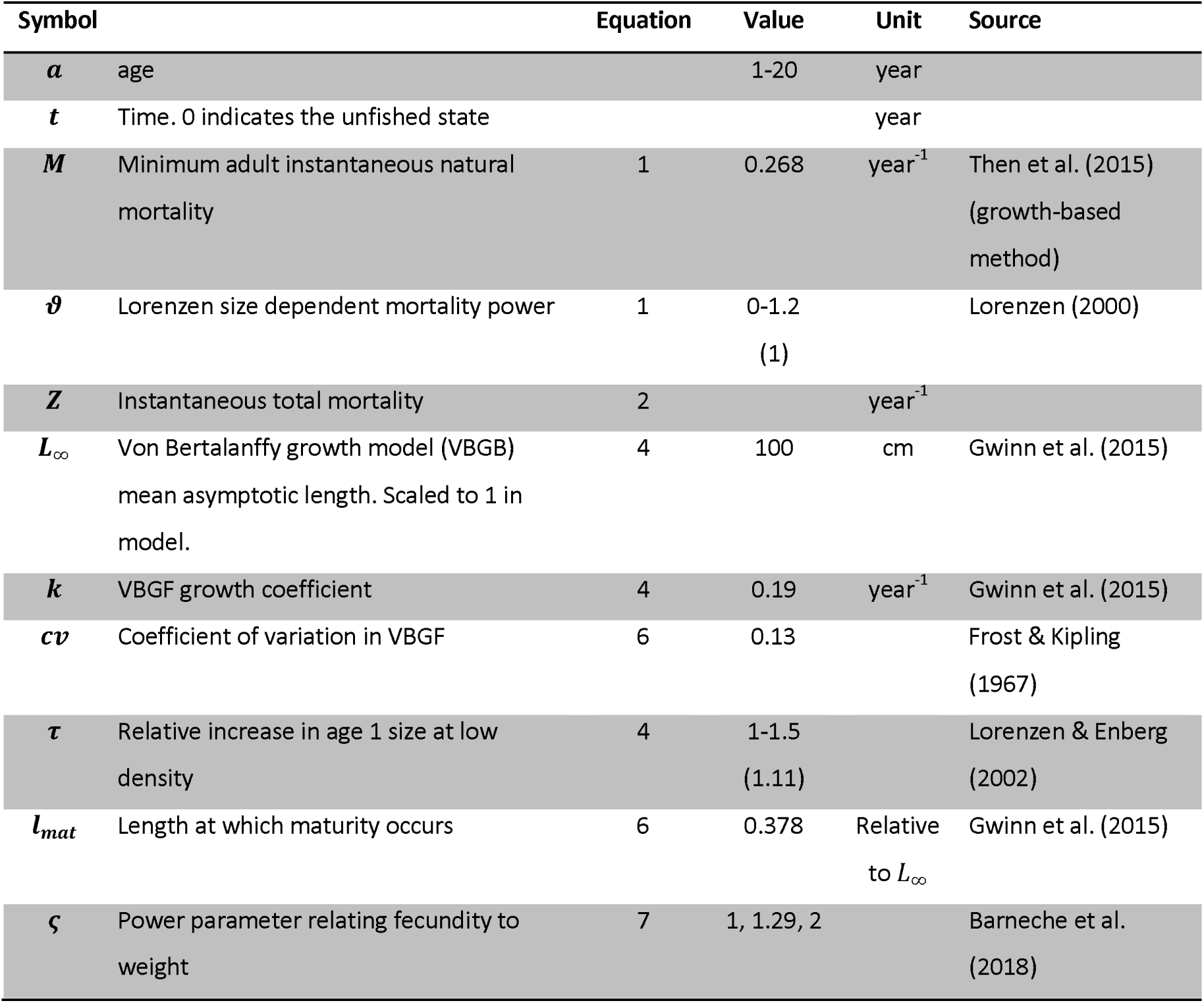
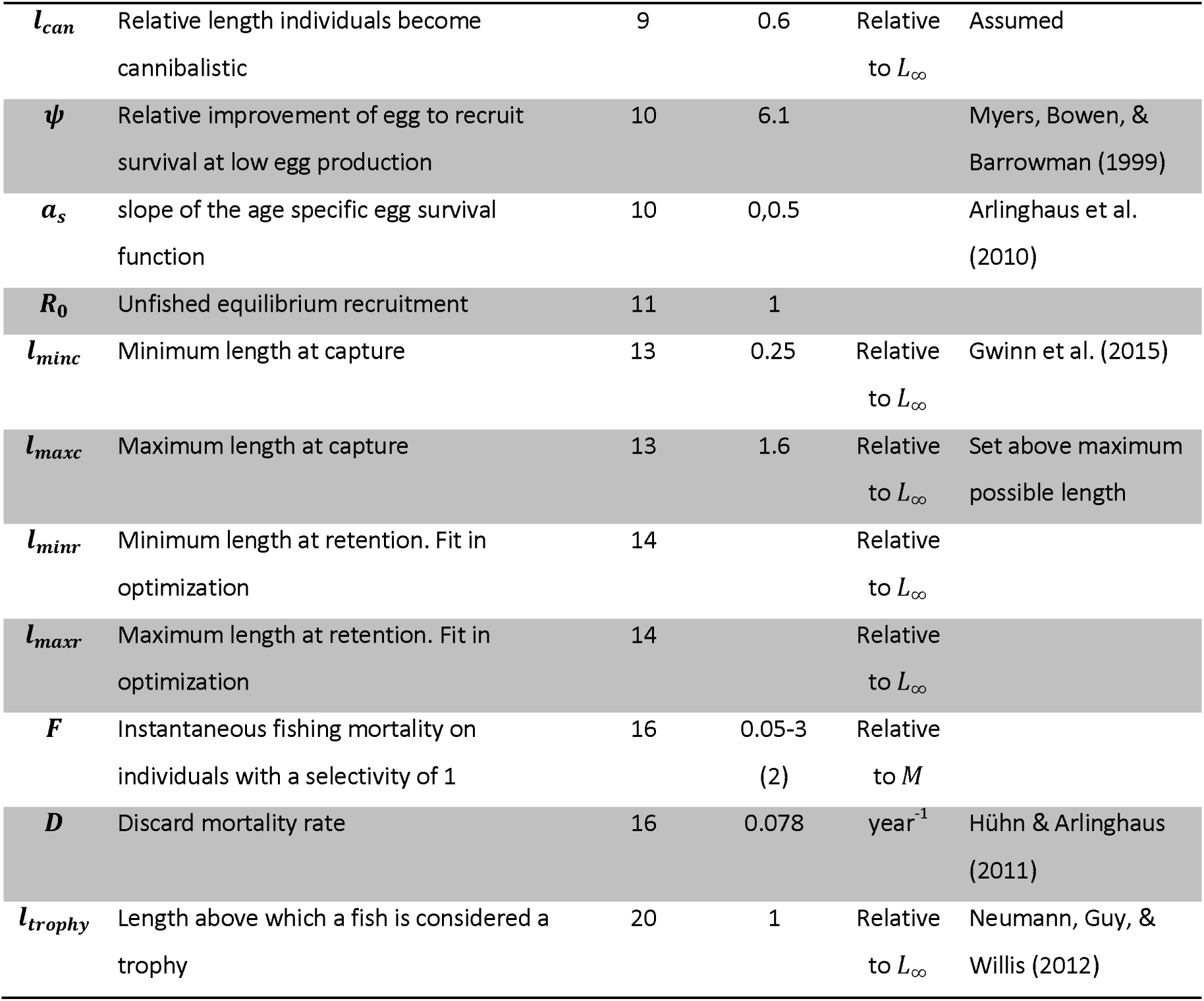
Parameters and parameter values, including units and where relevant sources. Values in parenthesis we used when the effects were held constant.

#### Mortality

Natural mortality (equation 1 in Table 1) was modeled as size dependent following the recommendations and empirical results of Lorenzen (2000, 2005). Accordingly, size-specific natural mortality was assumed a function of mortality at a reference length (Lorenzen, 2000). For this study, we chose a reference length of *L_m_,* the average asymptotic length of the fish in the population and calculated the reference mortality at *L*_∞_ following the growth-based method proposed by Then, Hoenig, Hall, & Hewitt (2015). To achieve a decline in natural mortality with increasing length of fish, the reference mortality at *L*_∞_ was multiplied by the ratio of the length at the reference age (*L*_∞_,) and length at a given age raised to a power ϑ (equation 1 in Table 1) as in Lorenzen (2000). In our model, we explored optimal size limits over a range of the strength of size dependency by varying ϑ in equation 1 from 0-1.2. The result was a decline in age-specific mortality with increasing size-at-age. Typical shapes of the size-dependent mortality are shown in Figure 2d. Across a range of species stocked at different sizes, Lorenzen (2000) found the most suitable value for the power ϑ to be 1 (called *c* in Lorenzen, 2000), and hence this value was used when size-based harvest policies were explored over a range of harvesting pressures.

**Figure 2.**
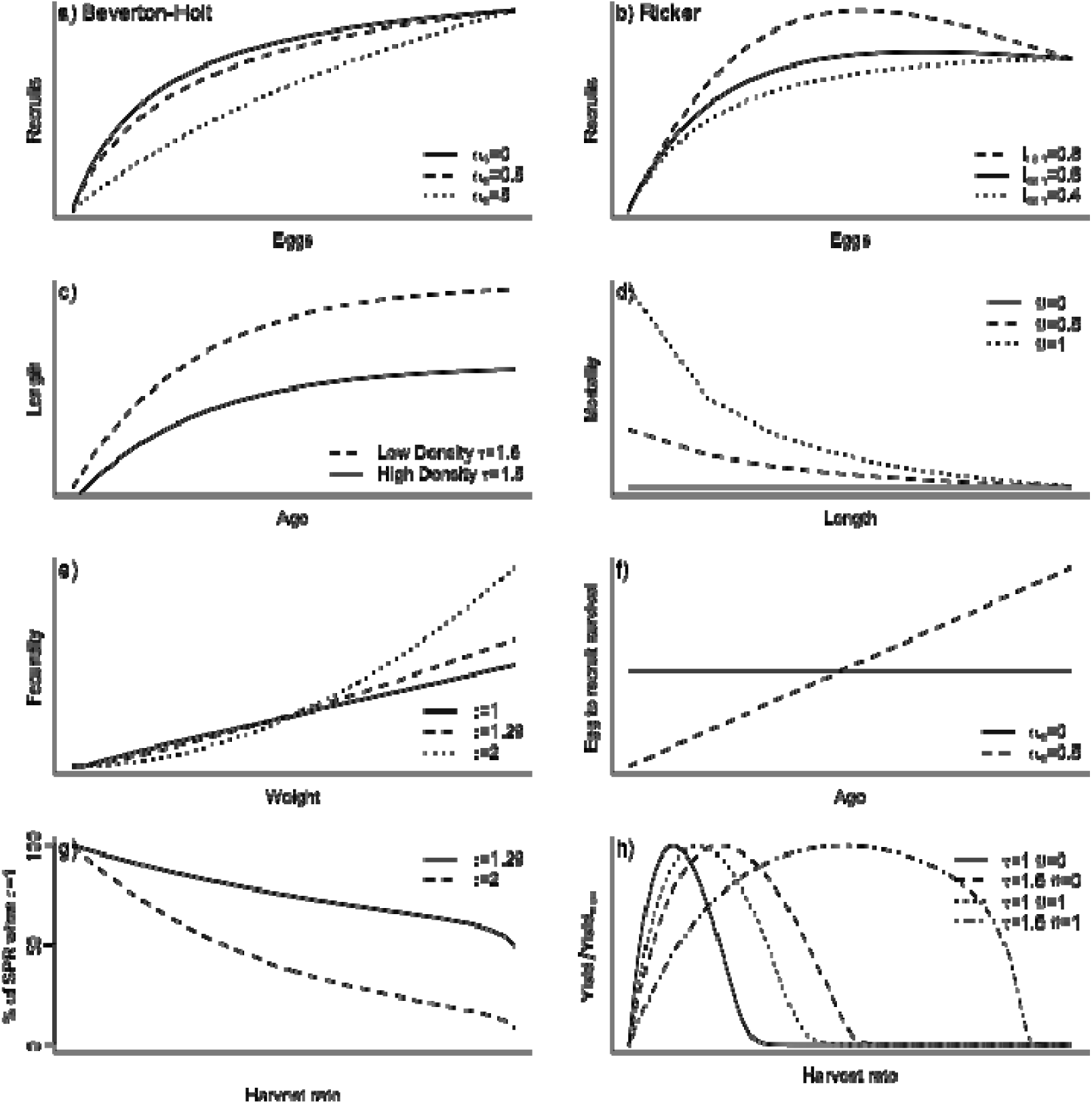
Key biological processes in the model and conceptual description of prototypical influences on biological and productivity rates. Notation follows Table 2.

#### Growth

Growth was assumed to follow the standard von Bertalanffy growth curve (von Bertalanffy, 1938) modeled using the Ford-Walford (Ford, 1933; Walford, 1946) linear equation (equation 4 in Table 1). Density dependent changes in growth were modeled assuming a relative increase (τ) in the size at age 1 as population density declined, where the population density effect was assumed proportional to “metabolic biomass” as the sum of the lengths squared in the population (rather than biomass) following recommendations by Walters & Post (1993). The relative increase in size with reduced population density was varied from 1­1.5 (for a possible effect on size-at-age, see Figure 2c). When size-based harvest policies were explored over a range of harvesting pressure a fixed value for density-dependence of growth of 1.11 (i.e., 11% decline in *L*_∞_ from low to maximum population density) was used as the average value reported by Lorenzen & Enberg (2002) for fish populations showing density dependence in growth.

Weight (i.e., mass at length) was modeled as the cube of length as is typical for fish (Walters & Martell, 2004). The combined effect of size dependent mortality and density dependent growth on relative yield can be seen conceptually in Figure 2h with both effects increasing the optimum harvest rate when judged as the harvest rate maximizing biomass yield.

#### Maturation and fecundity

Maturity-at-age was calculated as the proportion of individuals of a given age above a size threshold for maturation assuming variation in size-at-age was normally distributed around the von Bertalanffy growth curve with a constant coefficient of variation (equation 6 in Table 1) as is common in age-structured population models (Walters & Martell, 2004). Hyperallometry in fecundity with body weight was modeled as a power function of weight (equation 7 in Table 1) and scaled so that mean unfished eggs per recruit in the counterfactual isometric case (where absolute fecundity scaled linearly with body weight *ς* =1) was the same under the hyperallometric case (see Figure 2e). When hyperallometry in fecundity was explored relative to the isometric case, exponent values of 1.29 and 2 were used for the power *ς* of the mass-fecundity (egg number) scaling. An average value of the exponent of 1.29 across a wider range of marine fish species was reported in Barnache et al. (2018) for the weight-reproductive energy output scaling, while an average exponent of 1.18 was reported across species when only batch fecundity was considered. Specific for pike, Arlinghaus et al. (2010) previously used an exponent of 1.22, reflecting published work on the species, and Marshall et al. (unpublished data) presented an improved scaling of 1.89 for 26 species exhibiting repeat spawning. We thus choose a value of 1.29 to represent an extreme average case, and also explored a scaling exponent of 2 as the maximum exponent of the weight-fecundity relationship observed across several species by Barneche et al. (2018). The quite dramatic effect of hyperallometry on spawning potential ratio (SPR) can be seen in Figure 2h where the percent difference in SPR between the hyperallometric and isometric cases are shown.

#### Stock-recruitment and size-dependent maternal effects

Both Beverton-Holt (1957) and Ricker (1954) models for age-1 recruitment were modeled using Botsford incidence functions (Botsford, 1981a,b) following the methods for deriving equilibrium parameter values from Walters and Martell (2004) (equations 8-12 in Table 1, see Figure 2a,b). For both the Beverton-Holt and Ricker models, the maximum egg-to-recruit survival rate was calculated as the relative improvement in egg-to-recruit survival at low egg densities, known as the Goodyear compensation ratio *(ψ)* (Goodyear, 1980), times the egg to recruit survival rate in the unfished state (which reduces the reciprocal of eggs-per-recruit (0_eO_) in the unfished state) (equation 10a in Table 1). The Beverton-Holt scaling parameter (*β*), affecting how egg to age 1 survival changes as a function of total egg numbers, was modeled as a function of the unfished equilibrium number of recruits *(R_o_), ψ, φ_β0_* (equation 11a in Table 1). For the Ricker model, *β* was modeled as a function of *R_o_, ψ,* and the lifetime cannibalism impact per recruit (*ϕ*_canO_) (equation lib in Table 1). The proportion of individuals cannibalistic at each age was modeled similar to mortality, with a length threshold above which individuals were cannibalistic (Claessen, De Roos, & Persson, 2004). Survival to age 1 then varied as a function of the total number of cannibals in the population. As a higher proportion of individuals in the population became cannibalistic (a lowering of the size threshold), the Ricker model approached the Beverton-Holt form (see Figure 2b). The cannibalism size threshold set for simulations was 60% of *L*_∞_ so that the Ricker recruitment curve had a dome shape. This produced contrast between the two recruitment models. Sensitivity analysis of this and all other key parameters (Table 2) was completed to see whether the size limits optimizing biomass yield were elastic to parameter changes.

To explore the possible effect of increasing egg-to-recruit survival (size-dependent egg viability effect) with maternal age (and hence average size), maternal age-specific egg to age 1 survivals were modeled as a linear function of age (equation 10b in Table 1). If the slope *(a_s_)* of the age/survival relationship is specified (e.g. a 2.5 relative increase in egg-to-recruit survival from first time spawners at age 4 to age 9 females) then the intercept (a_0_) can be solved for to ensure that the maximum egg-to-recruit survival rate is equal that when no maternal age effect is assumed *(a_s_ =* 0) (see Figure 2f). The solution for *a*_0_ depends upon the age-specific proportional egg production (Ω_α0_) absent of harvest (equation 8 in Table 1), the lifetime proportional egg production per recruit (φ_Ω_), and the slope (a_s_) (rearrangement of equation 10b in Table 1). The viability effect of the maternal age relationship as we represented it can be seen in Figure 2a for the Beverton-Holt model: as the population declines through fishing and the older age classes are lost, the overall maximum egg-to-recruit survival declines resulting in a lower slope of the recruitment curve. Thus, as older individuals are removed the population becomes less productive because their proportional contribution to egg production (Ω_α_) declines resulting in a decline in the egg to age 1 survival rate 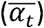 (equation 10c in Table 1). Note, however, that quite large relative differences in relative eggs-to-recruit survival are needed to cause large changes in the recruitment curve. When viability was modeled, a slope of 0.5 was used, representing a 2.5 fold higher egg quality of old pike relative to first time spawners as reported by Arlinghaus et al. (2010).

#### Fishing mortality and size-selectivity

Probability of capture and retention (equations 13 and 14 in Table 1) were modeled as the proportion of individual between the upper and lower size limits (representing size-selectivity) assuming variability in growth was normally distributed around the von Bertalanffy growth curve with a constant coefficient of variation. The minimum length at capture was set at 25% of the maximum *L_m_,* as in Gwinn et al. (2015). Upper and lower limits for the retention probability were selected in our model to optimize pre-defined management objectives. The difference between the capture and retention curves determined the proportion of individuals captured that were retained (equation 15 in Table 1), and individuals not retained were subjected to a release (i.e., discard or hooking) mortality (equation 16 in Table 1) as per Coggins et al. (2007). A mortality rate of 7.8% was used for the proportion not retained as this was the average hooking mortality reported in Hühn & Arlinghaus (2011) for pike that were angled.

Numbers at age were updated assuming continuous mortality (equation 17 in Table 1); a plus group for ages older that 20 years was not used because the mortality rates in our model resulted in age 20 being the maximum age (in agreement with reports in pike, Raat, 1988). Biomass and numbers harvested were calculated using the Baranov (1918) catch equation with appropriate accounting for capture and retention (equations 18 and 19 in Table 1). The proportion of individuals at each age greater that the average *L*_∞_ was used to calculate trophy catch (equation 20 in Table 1), assuming variability in growth was normally distributed around the von Bertalanffy growth curve with a constant coefficient of variation. This threshold comes from work by Neumann, Guy, & Willis. (2012) in recreational fisheries and personal knowledge that a pike of about 100 cm is considered a trophy in many cultures. Total trophy fish caught as well as numbers caught (not to be confused with harvest) were calculated using the Baranov catch equation (equations 21 and 22 in Table 1) and the appropriate age-specific capture probability.

#### Management objectives

Four management performance measures (biomass yield, numbers harvested, trophy catch and numbers captured) were explored. All four management measures were then combined when exploring a compromise management objective, as the sum of the natural logarithms of the values of each objective (equation 23 in Table 1) following Walters & Martell (2004). All base simulations were run at a value of the instantaneous fishing mortality rate *F =* 0.51 (representing an annual percentage harvest rate of 0.4, regularly reported for pike, Pierce, Tomcko, & Schupp et al., 1995), which together with the estimate of the instantaneous natural mortality rate Λ4=0.268 resulted in an estimate of *F/M* of about 1.9. At such fishing mortality rates, fish stocks are growth overfished (Walters & Martell, 2004; Zhou, Yin, Thorson, Smith, & Fuller, 2012) and may thus benefit from control of fishing mortality, implemented through size limits in our model.

#### Outline of analysis

We explored the size limits (minimum length limits or the combination of a maximum with a minimum length limit creating a harvest slot) that optimized either single objectives (biomass yield, numbers harvested, trophy catch or numbers caught) or a combined objective function representing the sum of the natural logarithm of each of the four objectives at equilibrium. We also explored other combinations of a biomass yield and one or more of the other objectives, but results were qualitatively identical and thus only the four-objective combined utility function is reported. Simulations (and results) proceeded in four steps:

1. First, optimal regulations for scenarios of varying density-dependence in growth and size-dependency in natural mortality were explored for the base case of a low size at first capture and high fishing mortality rate F/M of 1.9 that strongly exceeded the fishing mortality rate that would produce maximum sustained yield (MSY) (Walters & Martell, 2004; Zhou et al., 2012,). This “forced” the implementation of size limits to control unsustainable fishing mortality to meet management objectives. We also tracked the outcomes for each of the four management objectives at the best compromise regulations in the multi-objective utility function to study how well each objective performed at the best compromise.
2. Second, we fixed density-dependence in growth and size-dependency in natural mortality at average levels commonly reported for assessed fish stocks following Lorenzen (2000) and Lorenzen & Enberg (2002), and explored optimal size limits across a wide range of fishing mortality rates *F/M*.
3. The initial two steps explored the best-performing size limits. However, alternative size-limits may produce “pretty good” outcomes (Hilborn, 2007), which we defined as the size limit within 95% of the maximum value for each of the four objectives and the multi-objective utility function. To find such outcomes, we, third, searched for the regulation combinations that produced results within 95% of the optimum using a grid search, again fixing the density-dependence in growth and the size-dependency in natural mortality as per the typical values reported for fish stocks (see step 2).
4. Fourth, by turning again to an unsustainable maximum fishing mortality rate of roughly *F/M* of 1.9 as in step 1, we explored the effect of assumptions about hyperallometry in mass-fecundity scaling and a size-dependency in egg viability on optimal size limits. We chose commonly reported values (e.g., an exponent of the mass-fecundity relationship of 1.29, Barneche et al., 2018 and a 2.5 fold higher egg viability of the oldest relative to the first time spawners, Arlinghaus et al., 2010; Berkely et al., 2004b; Bravington et al., 2016; Venturelli et al., 2010) as well as extreme values for both processes (e.g., an hyperallometric exponent of 2) to examine generic patterns.
5. Finally, we explored model sensitivity to key model parameters (Table 2) by independently varying each parameter by ± 20% and examining the change in the optimal size limit for biomass yield. Model sensitivity also represented the possible applicability of the pike model to other life-histories (i.e., species or populations).

## Results

### 1. Optimal harvesting in light of single fisheries objectives

For the parameter set chosen and at fishing mortality strongly exceeding the minimum natural mortality rates of adults *M* (i.e., for F/M = 1.9), a minimum-size limit maximized equilibrium biomass yield independent of which stock-recruitment relationship was assumed (Figure 3). The optimal minimum-size limit became more liberal as the degree of density dependence and size-dependent mortality increased, with the effect of size-dependent natural mortality being somewhat stronger than the effect of density-dependent growth (Figure 3).

**Figure 3.**
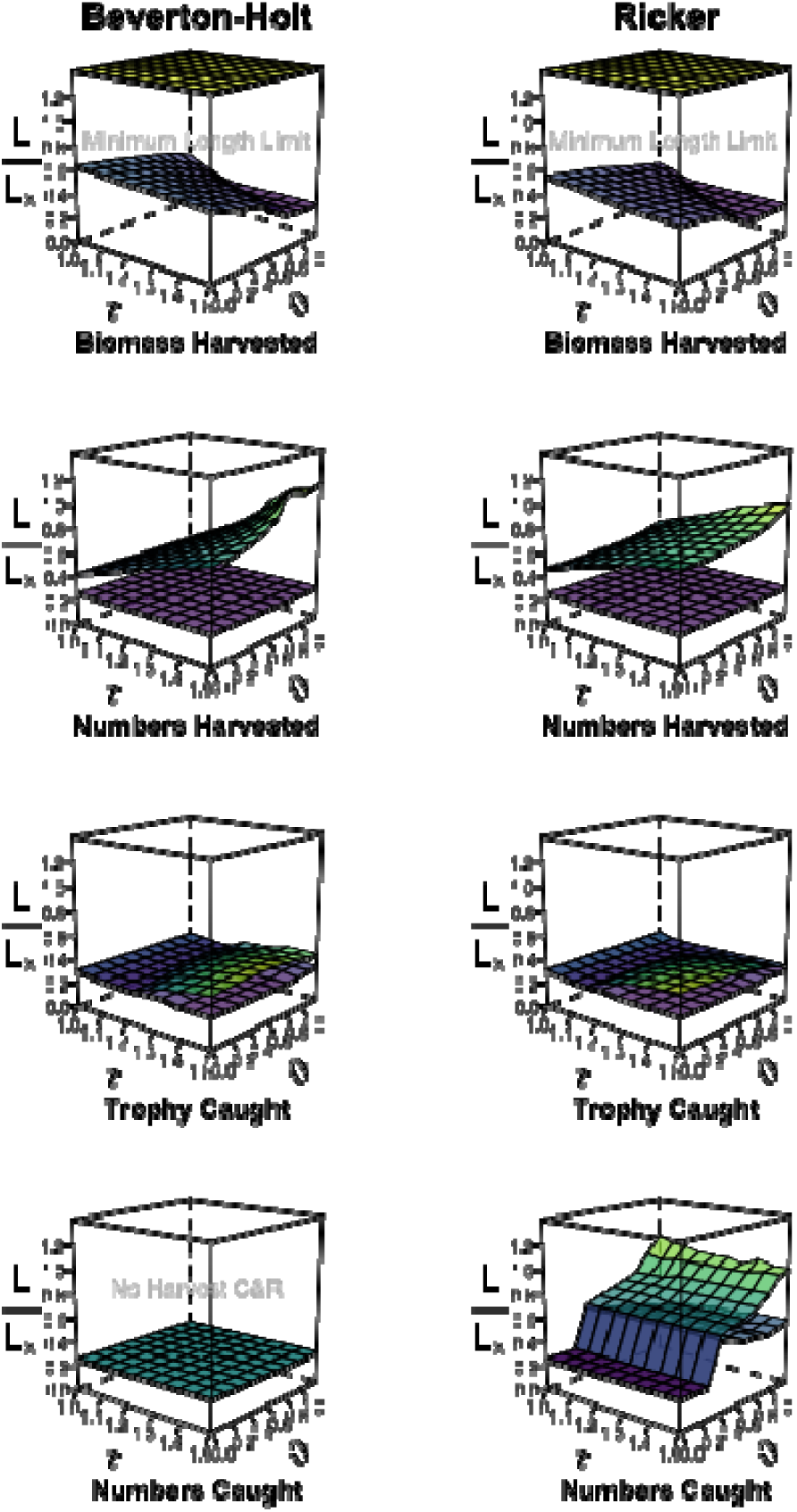
Impact of degree of density-dependent growth and size-dependent natural mortality on optimal size-selectivity for four fisheries objectives (size-selectivity is represented by where is the mean theoretical maximum length of the von Bertanlaffy growth model). The lower plane shows optimum minimum-length limits and the upper plane, when present, optimum maximum-size limit, together forming a harvest slot. The stronger the color gradient and the yellower the color of the upper plane, the larger are the differences between the minimum and the maximum-length limits. Note the maximum-size limit is only impactful when it is below 1.4, which is why a minimum-length limit is optimal for biomass yield. Whenever the planes diverge into two surfaces with different colors a harvest slot is the optimal harvest regulation. Results are shown for Beverton-Holt (left panels) and Ricker-type (right panels) stock recruitment across four management objectives, τ describes the degree of density-dependent growth, and ⍰ describes the degree of size-dependent natural mortality. C&R = catch-and-release indicating that the minimum and the maximum limits are identical, effectively creating a no harvest scenario.

Harvest slots, or more generally dome-shaped selectivity, optimized the numerical yield, independent of the stock-recruitment model (Figure 3). As before, as the degree of density-dependence in growth and size-dependency in mortality increased, the harvest slot became more liberal (i.e., widened), accommodating increased harvest. These effects were somewhat stronger under Beverton-Holt stock-recruitment compared to Ricker-type stock recruitment, which relates to differences by which survivorship relates to productivity (F_MSY_) in the two stock-recruitment models (see Martell et al., 2008 for details).

Harvest slots were also the optimal policy when maximizing the catch (rather than the harvest) of trophy fish, but this effect was only present when density-dependence in growth was strong, indicating that negative effects on growth at high density severely reduced the number of fish reaching trophy sizes supporting modest harvest through harvest slot-type regulations (Figure 3). By contrast, given weak density-dependence in growth, catch-and-release (i.e., a zero harvest policy) was identified as the optimal regulation for trophy catch under both Ricker and Beverton-Holt stock recruitment. When a harvest slot was optimal at strong density-dependence in growth, strong size-dependent mortality narrowed the optimal width of the harvest slot for both stock-recruitment curves - an effect somewhat more pronounced with Ricker recruitment.

The numbers captured were maximized through a total catch-and-release policy in the case of Beverton-Holt recruitment. That is, zero harvest was optimal for keeping the abundance at maximal levels, in turn maximizing the catch per unit effort. The situation was very different under Ricker stock-recruitment. Here, at low levels of size-dependent mortality, a total catch-and-release policy maximized catch rates, but at high levels of size-dependent mortality a harvest slot was optimal for achieving high catch numbers. The reason was that the resulting mortality shifted the recruitment to the maximum point in the Ricker curve, thereby increasing abundance and hence catch rates.

### 2. Optimal compromise harvest regulations in light of multiple objectives

Harvest slots turned out to be the optimal harvest regulation when considering four fisheries objectives - biomass yield, numerical harvest, trophy catch and catch rate - jointly in the log-utility function. This result was independent of the stock-recruitment relationship and was also independent of density-dependence in growth and size-dependency in natural mortality (Figure 4). As before, as the strength of density-dependence and size-dependence increased, the harvest slot widened, allowing more intensive harvesting. Ricker-type stock recruitment allowed more aggressive harvesting compared to the Beverton-Holt case because exploitation reduced the degree of cannibalistic control through the larger size classes on recruits.

**Figure 4.**
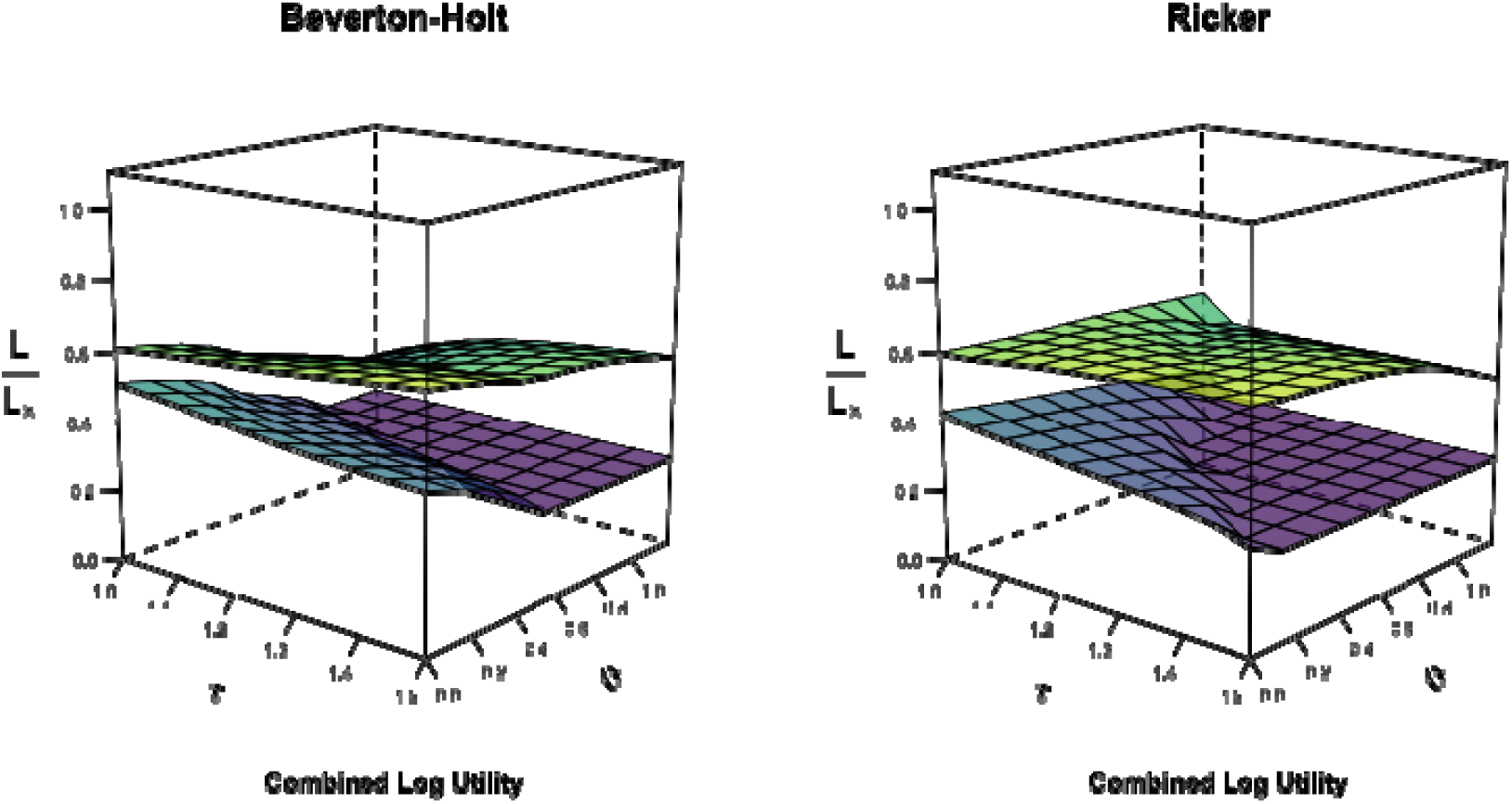
Optimal compromise harvest regulation when using a combined log utility function to integrate the four management performance measures shown in Figure 3. The impact of degree of density-dependent growth and size-dependent natural mortality on optimal size-selectivity is represented by where is the mean theoretical maximum length of the von Bertanlaffy growth model. The lower plane shows the optimum minimum-length limits and the upper plane, when present, the optimum maximum-size limit. The stronger the color gradient and the yellower the color of the upper plane, the larger are the differences between the minimum and the maximum-length limit. Whenever the planes diverge into two surfaces with different colors a harvest slot is the optimal harvest regulation. Results are shown for Beverton-Holt (left panels) and Ricker-type (right panels) stock recruitment, τ describes the degree of density-dependent growth, and 0 describes the degree of size-dependent natural mortality.

The fisheries outcomes using a harvest slot as a compromise regulation achieved pretty good outcomes for each of the four performance measures relative to the maximum possible outcome for each of the objectives (Figure 5). As characteristic for a compromise, no single measure was maximized under the compromise harvest slot. Yet, in all cases, the average outcomes were > 50% of the maximum possible, which we interpret as “pretty good”. The compromise harvest regulation achieved over 90% of the maximum possible harvest numbers under both the Ricker and Beverton-Holt scenarios. The Beverton-Holt compromise regulation also achieved among 60 and 70% of the maximum in the catch of trophies and the numbers harvested, and roughly 55% of the maximum biomass yield, on average. The Ricker compromise harvest slot performed better at the biomass yield level, with a value closer to 70% and poorer than the Beverton-Holt case on the trophy catch (only slightly above 50% of the global maximum).

**Figure 5.**
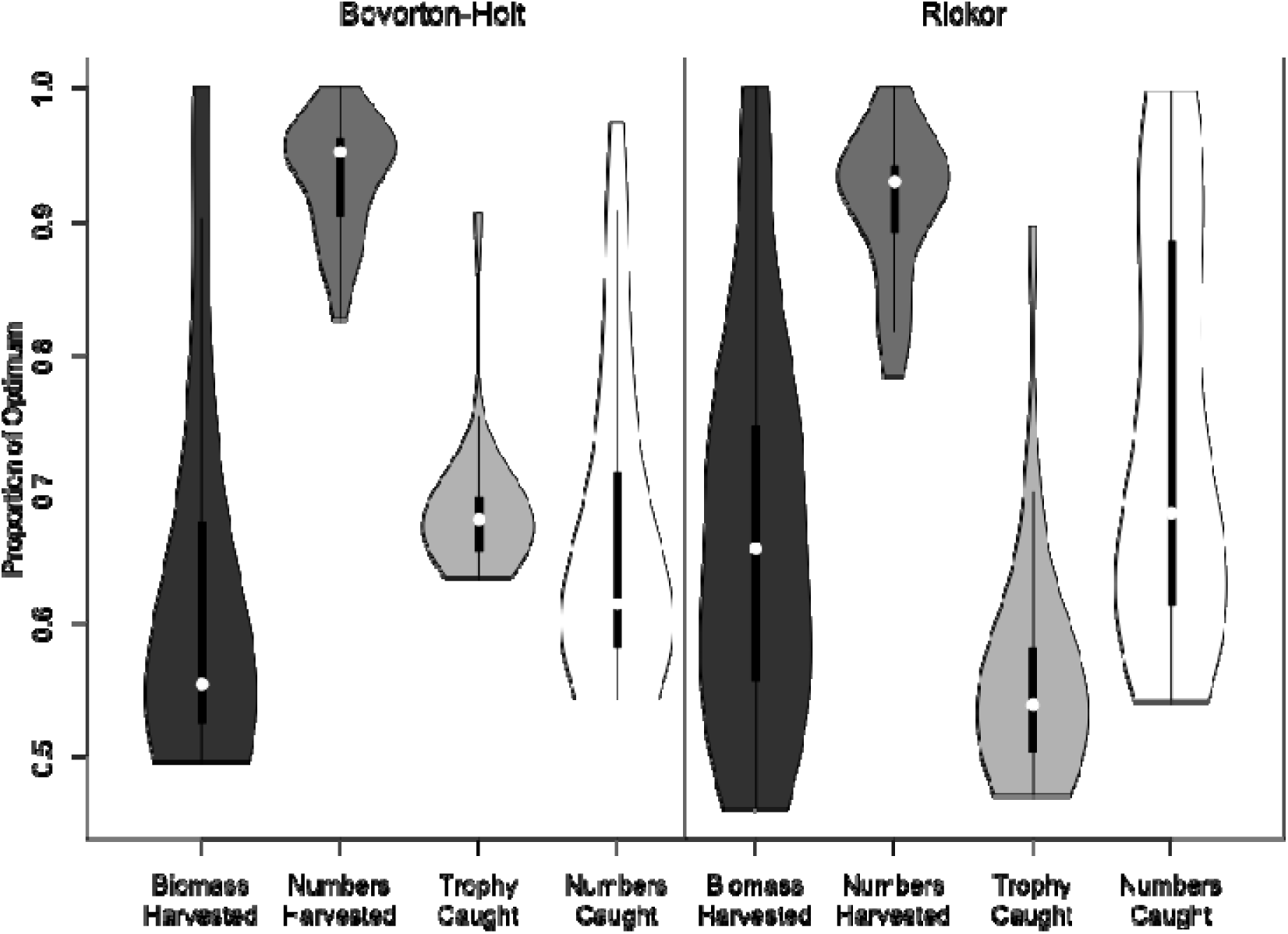
Outcomes at the optimal compromise harvest slot (from Figure 4) for the entire set of ecological scenarios of density-dependence in growth and size-dependence in natural mortality shown in Figure 3 across four objectives. We define pretty good outcomes when each objective is larger than 50% of the maximum theoretically possible for the parameter set that was chosen. All objectives met that criterion.

### 3. Optimal harvest regulations with varying fishing pressure

Fixing density-dependence in growth and size-dependent mortality at parameter values commonly reported for exploited fish stocks allowed systematic examination of how the optimal harvest policies varied with total fishing pressure. At equilibrium, minimum-length limits were the optimal harvest regulation for maximizing biomass yield across a large fishing pressure gradient (Figure 6). However, particularly under Ricker-stock recruitment, harvest slots started to appear as optimal for biomass yield when fishing pressure was high.

**Figure 6.**
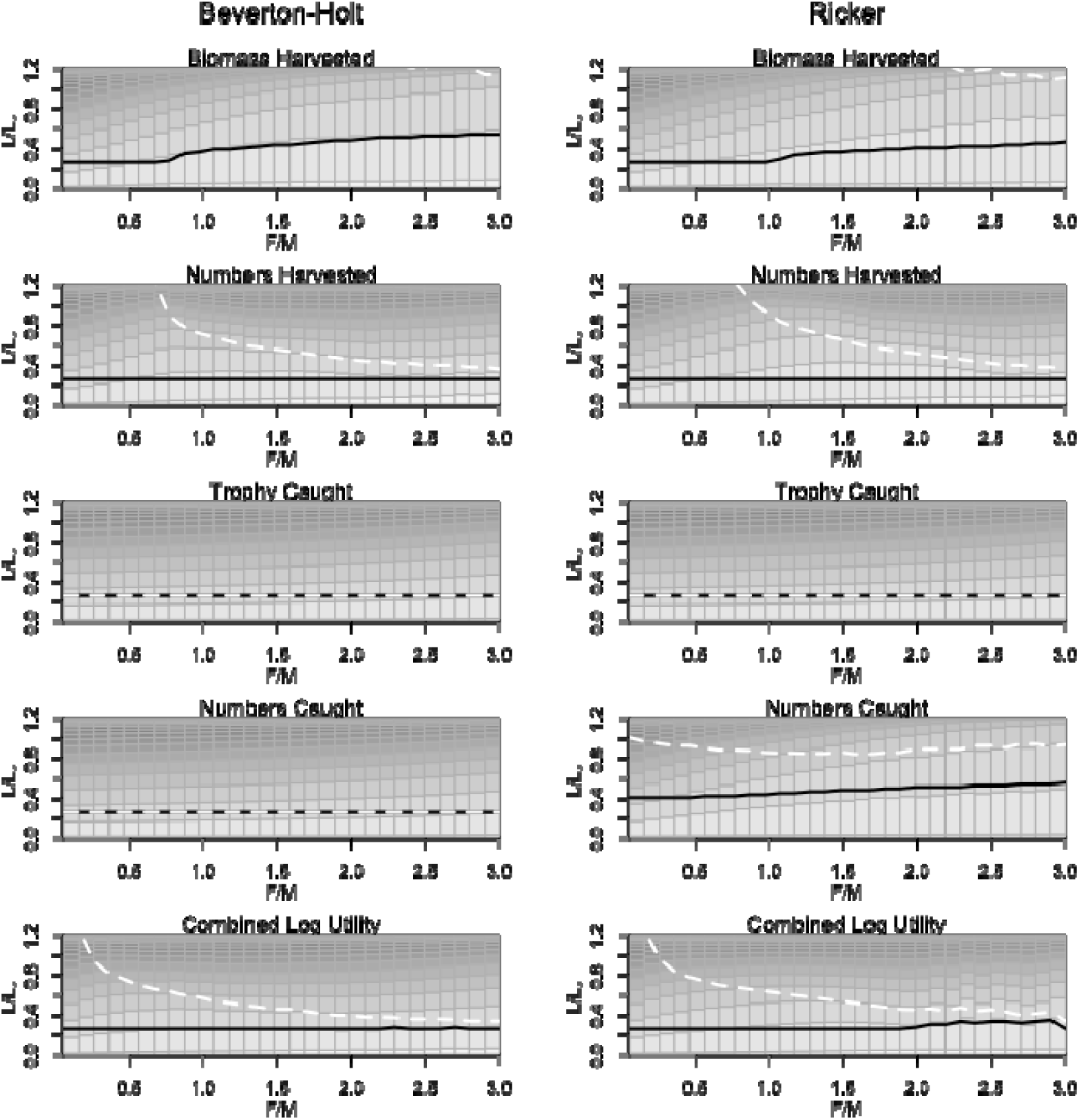
Optimal harvest regulations by management objective, for four individual objectives and a combined objective function, along a gradient of fishing pressure F/M. The white broken line indicates the upper limit of the harvest window, while the solid black line indicates the lower minimum-length limit. The length limits are expressed relative to, the maximum mean length of the von Bertanlaffy growth model. The bars in the surface for each level of maximum fishing mortality indicate the egg contribution by different age classes for a given fishing pressure. Younger ages are shown in lighter grey, older ages in darker grey. All simulations were done with an average degree of density-dependent growth and size-dependent natural mortality.

Harvest slots started to constitute the optimal regulation for maximizing harvest numbers at fishing mortality rates of *F* exceeding 0.5 *M.* At lower fishing mortality rates, the minimum length limits were equivalent to the minimum capture size (25% of mean L_æ_), indicating no regulation at all was needed at low fishing mortality rates.

The catch of trophy fish was optimized by a zero harvest policy, independent of the fishing pressure (Figure 6). Similarly, a no harvest policy was revealed as the optimal regulation to maximize catch in numbers under Beverton-Holt stock-recruitment across the fishing pressure gradient. However, when the stock-recruitment function followed a Ricker form, catch numbers (and hence abundance) were maximized with a harvest slot, independent of the fishing pressure (Figure 6).

Independent of the stock-recruitment curve and the local fishing mortality, harvest slots best compromised among the four fisheries-management objectives. With increasing fishing pressure, the width of the optimal harvest slot narrowed, indicating constrained harvesting with higher fishing pressure.

The different optimal harvest regulations had characteristic impacts on the age structure of the stock and the relative contribution of different age classes to total egg production (Figure 6). This is most clearly seen when comparing the distribution of eggs by age class under the optimal regulations for biomass yield relative to those that optimize numerical harvest. Under the biomass maximization objective, a strong juvenescence effect is visible, and the bulk of the eggs were produced by young age classes. By contrast, the implementation of harvest slots to maximize numbers harvested reversed the juvenescence effect, and the relative contribution or older ages to egg production increased as the harvest slot narrowed (Figure 6). Essentially, with a harvest slot the older age classes served as the reservoir to produce new recruits, which were intensively harvested as long as they continued to grow into the slot. However, under Ricker-type stock recruitment the rather wide harvest slot needed to produce maximum recruitment and hence maximum catch rates resulted in a strong juvenescence effect, indicating that the large cannibals were the ones that were removed if the aim was to increase catch rates. By contrast, the compromise regulation achieved a very balanced age structure, represented by a relatively even contribution of different age classes to the total eggs, mirroring the pattern under no harvest (see the trophy panel in Figure 6).

### 4. Size-limits that produce “pretty close” outcomes within 95% of the optimal

Several regulation combinations produced outcomes that were equivalent or “pretty close” to each other. We examined this pattern by searching for all size limits that produced outcomes within 95% of the optimum (Figure 7, the optimum size limits for comparision are shown in Figures 3 to 6). In many cases harvest slots produced similarly good outcomes as minimum-length limits alone (Figure 7). For example, a minimum-length limit was found to be the single optimal regulation to maximize biomass yield for a wide range of fishing pressures for both Beverton-Holt and Ricker-type stock recruitment (Figure 6). However, implementing a modestly large maximum-size limit in addition a small minimum-length limit produced biomass yield within 95% of the maximum possible over a wide range of fishing pressures (Figure 7). For example, at a fishing pressure of F/M = 1 minimum length limits ranging from 0.25 to slightly above 0.4 of L_æ_ would in combination with a maximum-size limit of > 0.8 of L_æ_ create similarly good outcomes for yield within 95% of the maximum.

**Figure 7.**
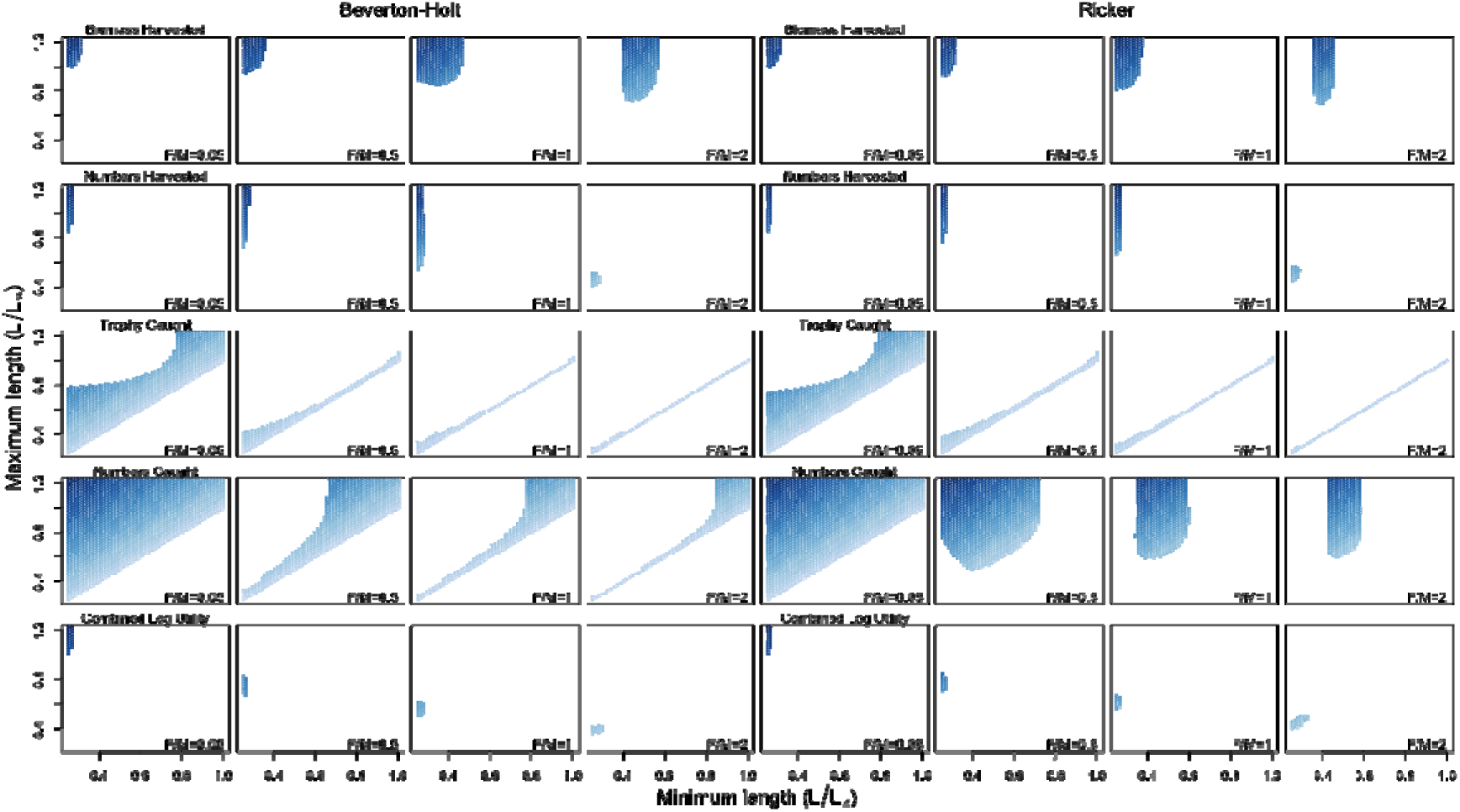
Minimum and maximum length limits expressed as, where the combination indicates a harvest slot that produces at least 95% of the objective’s maximum possible across different fishing pressures (expressed as *F/M).* Four objectives across either a Beverton-Holt (left panels) or a Ricker stock recruitment (right panels) relationship are shown. Darker blue indicates larger differences between the maximum and minimum limits. Whenever a bar, rather than a point, is shown it indicates that various combinations produce pretty close outcomes. A diagonal line shown for some trophy catch cases represents the 1:1 line where minimum and maximum-length limits are equal and thus catch-and-release (no targeted harvest) is the optimal policy. All simulations were done at an average degree of density-dependent growth and size-dependent natural mortality, commonly reported in the fish ecological literature.

Optimizing the harvest numbers would require keeping the minimum-length limit low, confined to about 0.25 of *L*_∞_ across all fishing pressures, but adding a maximum size limit of a wide range would create similar good outcomes for harvest numbers. The higher the fishing pressure, the lower the maximum-size limit would need to be to achieve pretty close results, but this opportunity of flexible upper limits with similar outcomes vanished to a confined harvest slot at *F/M* of about 2 under both Beverton-Holt and Ricker (see also Figures 3 and 6 showing a scenario for F/M of about 1.9).

Optimum outcomes for trophy catch depended on catch-and-release (indicated by minimum and maximum size-limits being identical, visualized along the diagonal in Figure 7) unless the fishing pressures was low. At low fishing pressure the impact of undesired discard mortality vanished, in which case a wide range of harvest slots and associated combinations of minimum and maximum length limits were conceivable to produce equally good outcomes for trophy catch.

Similarly, for numbers captured, the zero harvest optimal policy could be substituted by a wide range of size limits achieving “pretty good” results at low fishing pressure for both recruitment models. As fishing pressure increased these options reduced to retention of only the largest individuals in the Beverton-Holt model and the emergence of a harvest slot limit and low size limit in the Ricker model.

When seeking the best compromise regulation very limited flexibility was predicted. Under basically all cases of fishing morality, a confined harvest slot was found to be best, with the upper limit decreasing as the fishing pressure increased. The minimum-length limit of this optimal compromise was consistently small and offered very limited leverage if the goal was to achieve within 95% of the maximum possible outcome for the compromise.

### 5. Impact of hyperallometry in fecundity and size-dependent egg viability on optimal harvest regulations

The introduction of size-dependent maternal effects, both in terms of egg production (fecundity increasing non-linearly with mass with an exponent of either 1.29 or 2) and egg viability (where the largest fish produce eggs that are 2.5 times more viable than the first­time spawners) had overall modest effects on the optimal size limits for biomass harvested, numbers harvested, trophy catch and the compromise regulation, with effects being somewhat stronger under Ricker recruitment for biomass yield and a bit less pronounced on harvest numbers compared to the Beverton and Holt case (Figure 8). Introducing a viability benefit for eggs spawned from large spawners had almost no impact on optimal regulations, independent of assumptions with our without additional hyperallmometry in fecundity. Stronger effects were seen for assumptions of hyperallometry in fecundity, but effects were only pronounced for some objectives when the assumption was made the fecundity scaled with body mass with an exponent of 2. Under this assumption, a harvest slot (Figure 8) rather than a minimum-length limit (Figure 6) was found optimal for biomass yield under both stock-recruitment models at fishing pressures *F/M* above 0.5-1. Similarly, the harvest slot option appeared to be optimal at smaller fishing pressures under hyperallmometry than under isometry for harvest numbers, particularly for Ricker stock-recruitment, where the slot narrowed compared to assumptions of isometry in fecundity. Size-dependent maternal effects particularly affected the optimal regulation on catch rates under Ricker stock-recruitment, with the harvest slot that produced highest catch rates narrowing with non­linearly increasing fecundity with mass and less so with assumptions of higher egg viability for large individuals.

**Figure 8.**
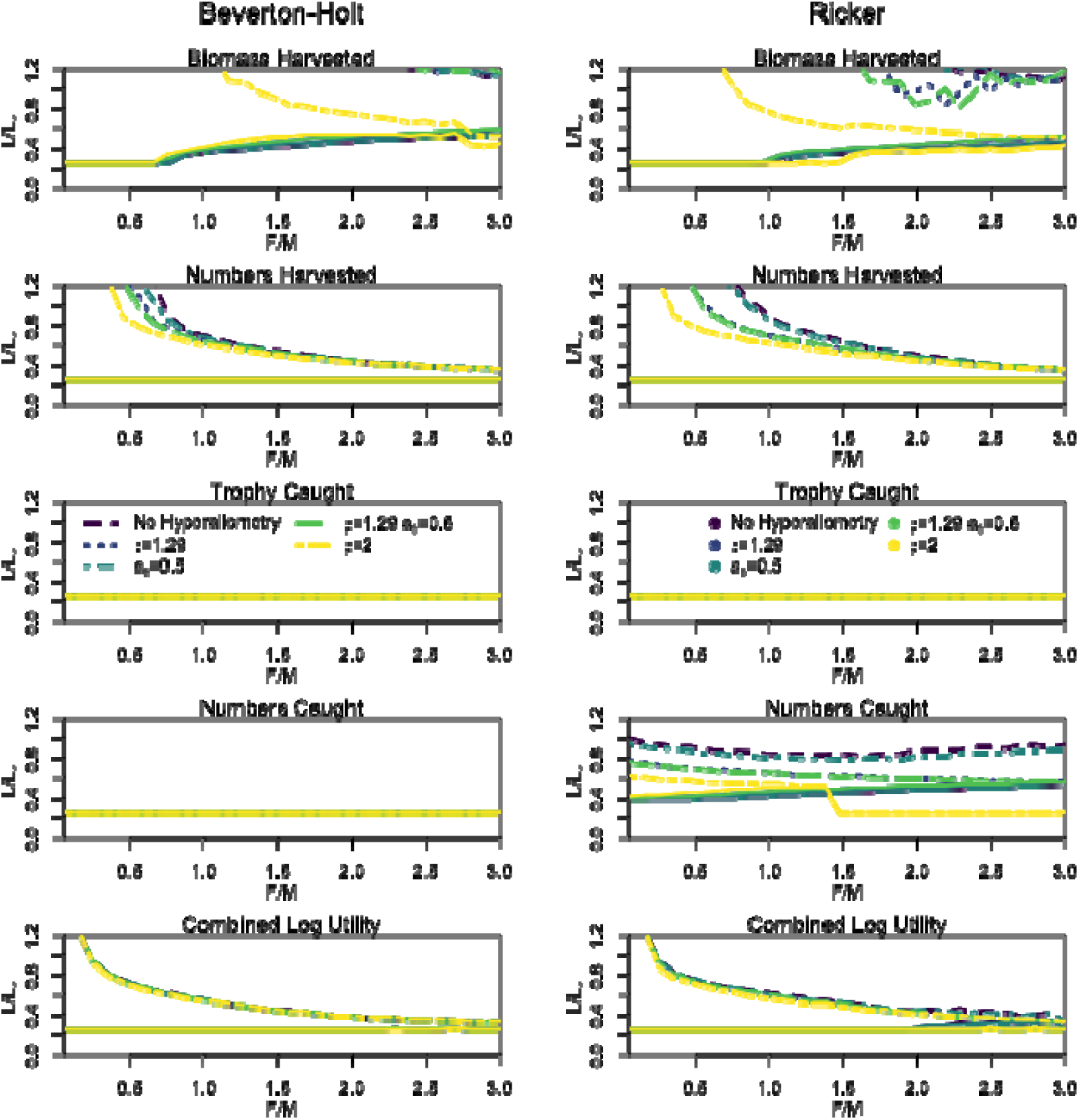
Optimal harvest regulations by management objective and for a combined utility function along a gradient of fishing pressure when hyperallometry in fecundity and size-dependent egg viabilities were assumed alone or in combination. Colored solid lines indicate the minimum-length limit and same colored broken lines indicate the upper limit of the harvest slot whenever it is considered optimal.

### 6. Robustness of model predictions to parameter uncertainty

The sensitivity of the model to the main parameters was evaluated by analyzing the percent change in the optimal lower size limit under a biomass harvest maximization policy for both Beverton-Holt and Ricker recruitment with a 20% change in each parameter (Table 3). As expected, the optimum lower size limit was most sensitive to parameters impacting mortality and growth, with the minimum adult instantaneous mortality () and the relative change in size at age 1 at low population density (τ) being the most important parameters. Increases in *M* resulted in declines in the minimum length limit. Increases ίητ also resulted in reduction in the size limits as a result of increases in the proportion of individuals reaching maturity at younger ages. Both the Lorenzen size-dependent mortality power (i9) and the von Bertalanffy growth coefficient *(k)* produced moderate sensitivity. An increase in *ϋ* resulted in declines in the minimum length due to increases in the average mortality rate in the population. Increases in *k* resulted in declines in the minimum length limit due to increasing average mortality-at-age, which had a stronger effect than decreases in the proportion of individuals maturing. Changes in other parameters resulted in low sensitivity. Effects were similar for both recruitment models.

**Table 3.** Sensitivity of the optimum (for biomass yield) minimum length limit to model parameters, with the percent change in the lower length limit for the biomass maximization management option as a result of a 20% in the parameter. Values in bold indicate more important (i.e., sensitive) parameters with a greater than 20% change to a changing input parameter. * Density dependent growth scaler could only be reduced to 1 and the results show only a 10% reduction. Weight power scaling on fecundity and maternal age effect on viability were only increased as base runs were at the minimum possible values. Notation follows Table 2.

| Parameter | Beverton-Holt |  | Ricker |  |
| --- | --- | --- | --- | --- |
|  | -20% | +20% | -20% | +20% |
| $M$ | **28.0%** | **-41.2%** | **15.3%** | **-31.0%** |
| $\vartheta$ | 17.5% | -16.9% | 7.8% | -18.3% |
| $k$ | -14.0% | 14.2% | -16.3% | 8.7% |
| $cv$ | -0.3% | 1.1% | 0.4% | -1.2% |
| $\tau^*$ | **21.9%** | **-26.8%** | **6.6%** | **-31.0%** |
| $l_{mat}$ | -4.9% | 8.2% | -14.1% | 7.2% |
| $\varsigma$ | - | 6.0% | - | 4.4% |
| $l_{can}$ | - | - | 1.3% | -1.2% |
| $\psi$ | 6.9% | -3.0% | 5.9% | -4.7% |
| $a_s$ | - | 0.3% | - | 0.3% |
| $l_{minc}$ | 0.4% | -0.2% | 0.2% | -0.1% |
| **D** | 1.3% | -0.3% | 0.6% | -0.4% |

## Discussion

Common sense and decades of empirical and theoretical work suggests that letting fish spawn at least once usually safeguards recruitment, thereby avoiding recruitment overfishing (Myers & Mertz, 1998). Moreover, letting the fish growth until a cohort reaches its maximum biomass before harvest constitutes a suitable approach to achieve high biomass yields (Froese et al., 2016). A minimum-length limit set well above size at maturation is thus predicted to maximize yields (Allen, 1953; Ayllón et al., 2019; Beverton & Holt, 1957; Clark et al., 1980; Dunning, Ross, & Gladden, 1982; Gwinn et al., 2015; Jensen, 1981; Lenker et al., 2016; Prince & Hordyk, 2019; Reed, 1980; Ricker, 1945; Saila, 1956; Van Gemert & Anderson, 2019). In support of this classical perspective, our model similarly predicts that minimum-length limits are a suitable harvest regulation if the aim is to achieve high biomass yields.

We add to this established literature that the biomass-maximizing effects of minimum-length limits is largely independent of the degree of density-dependence in growth, size-dependent mortality and size-dependent fecundity (Figures 3,6). However, situations change when hyperallometry in fecundity is strong. Under such scenario, biomass yield is predicted to be maximized with harvest slots at high fishing pressures by offering some protection to the highly fecund large fishes (Figure 8). Our model also showed that under assumption of isometry in fecundity adding a maximum-size limit on top of a suitable minimum-length limit, thereby creating a harvest slot, does not cause a substantial decrease in yield and can produce within 95% of the biomass yield promised by a minimum-length limit. Consistently, we found harvest slots to constitute the optimal regulation for numerical yield at moderate to high fishing mortality rates, for trophy catch under strong density-dependence in growth and for catch numbers under strong size-dependent mortality and Ricker type stock recruitment. Overall, optimal policies need to be judged against predefined objectives and thus cannot easily be generalized.

We also found a harvest slot to consistently constitute the best-performing regulation when integrating four typical fisheries objectives. In fact, a harvest slot always emerged when at least one other fisheries objective or performance measure was added to a biomass yield objective, particularly when catch- or catch size-based objectives were in place that prefer lower fishing mortality rates than those that produce MSY and reduce equilibrium biomass. This finding extends the work by Gwinn et al. (2015) to a much more general case because we examined variation in stock-recruitment relationships (representing density dependent juvenile mortality), density-dependent growth and size-dependency in mortality and fecundity/egg viability, thereby simulating a large family of ecological and species-specific population processes that the Gwinn et al. (2015) model lacks. Moreover, similar to Gwinn et al. (2015) we found our results to be very robust to variation in life-history traits (thereby representing other life-history prototypes or species than the pike). In fact, while variation in life-history traits (e.g., growth rate) will affect the overall productivity (e.g., yield or F_MSY_) of the stock (e.g., Martell et al., 2008) and thus the exact configuration of an optimal policy (e.g., in terms of width of the optimal harvest slot), the optimal policy per se (i.e., whether a minimum-length limit or a harvest slot is optimal) is unlikely to change much. One exception identified in our work is the presence of extreme hyperallometry in fecundity which may shift the optimal policy for biomass yield from a minimum-length limit to a harvest slot at moderate to high fishing pressures. Therefore, in light of the robustness of our results to life-history variation we content that in fisheries where multiple objectives are to be achieved jointly that encompass both extraction and catch-related objectives, harvest slots or other types of dome-shaped selectivity, i.e., selectivity patterns that protect both immature and very large mature fish may constitute superior harvest strategies to the standard minimum-length limit-type regulation for a wide range of species (see also Gwinn et al., 2015).

### Impact of stock-recruitment

Our results were largely robust to assumptions about the underlying stock-recruitment relationship. Assuming different types of stock-recruit relationships can be interpreted to represent differences in reproductive biology across a large famility of species, and the robustness of our results imply some level of generality of our findings to apply broadly beyond pike (and other cannibalistic species). However, some exceptions are worth noting that appeared when Ricker-type recruitment and hence an impact of cannibalistic intraspecific control was assumed. In particular, for catch numbers (a surrogate for catch rate) under a Ricker model a harvest slot rather than a zero harvest policy was found to be optimal. The reason can be found in the so-called overcompensatory feature of the Ricker stock-recruitment model (Ricker, 1954). When the abundance of large cannibals increases, these individuals may strongly reduce recruitment through inter-cohort predation (Persson et al., 2006). When the cannibals are removed, either through some modest harvest or through discard mortality at high fishing pressure, the recruitment initially rises, as for example found in pike (Sharma & Borgstrpm, 2008, see also Figure 2b). This boost in recruitment maximizes abundance and hence catch rates, at the potential conservation cost of truncation in size and age structure. Overcompensation as predicted from cannibalism is the key difference between the Beverton Holt and the Ricker model, which explains why some modest harvesting is needed to maximize a property such as numbers captured under a Ricker model. Typically, maximized abundance, and hence maximized catch rate, is associated with zero harvest and unexploited conditions (Beverton & Holt, 1957; Hilborn, 2007). Worldwide, most stocks for which data is available follow a Beverton-Holt stock-recruitment relationship, but there are 17% of global stocks for which data are available that show Ricker recruitment (Szuwalski et al., 2015). In particular strongly piscivorous, and by the same token cannibalistic marine and freshwater species, such as pike-perch *(Sander lucioperca,* Grôger, Winkler, & Rountree, 2007), walleye *(Sander vitreus)* (Zhao, Kocovsky, & Madenjian, 2013), pike (Edeline et al., 2008) or cod (Sguotti et al. 2019) show evidence for Ricker-recruitment. Hence, based on our model, in top predatory species modest harvest is recommended even when the goal is to maximize catch rate or catch of trophies, as some harvest releases the remaining fish from cannibalistic control, increasing recruitment into the fishery and growth of fish to reach memorable, trophy sizes.

### Density-dependent growth and size-dependent mortality

*We* show that population resilience substantially increased with density-dependent growth and with size-dependent mortality. The increased resiliency to harvest with increased size-dependent mortality can be explained by the increase in average natural mortality rate of the exploited stock when size-dependent mortality is present compared with the situation when it is not. Increased natural mortality rate means the stock turns over faster and fisheries can take fish that would otherwise die naturally, increasing sustainable harvest rates (Lester, Shuter, Venturelli, & Nadeau, 2014; Martell, Pine, & Walters, 2008; Zhao et al., 2013).

The mechanism for increased resilience to harvest caused by density-dependent growth is different and more complex. With density-dependent growth, the fish grow faster when they are exploited, which increases the biomass gain per unit time and thus the stock becomes more productive. Importantly, growth plasticity means that exploited fish reach the maturation size threshold earlier and produce more eggs at a given age. Reductions in age at maturation increase the compensatory reserve and allow more intensive harvesting. There is a third effect of density-dependent growth, which is to reduce the average natural mortality rate when there is size-dependent mortality in addition to density-dependent growth. All else being equal, the effect of decreasing average natural mortality alone would reduce resiliency to harvest, but in our model the compensatory growth and maturity effects overcompensate this effect, increasing resiliency to harvest under density-dependent growth.

Our model showed that the optimal size-based harvest regulations were largely robust to assumptions of density-dependent growth and size-dependent mortality. Yet, the exact configuration of the optimal size limit changed as assumptions about these properties changed. We found that when density-dependence was strong, maximizing trophy catch necessitated some modest biomass removal, or discard mortality through high effort, to release the remaining fish from density-control, in turn fostering growth into trophy sizes. The concern that high abundances, for example caused by anglers engaging in total voluntary catch-and-release in selected fisheries, may jeopardize trophy catches is frequently expressed for recreationally important trophy species, such as muskellunge (Gilbert & Sass, 2016). Our model supports these concerns and offers a solution. Clearly, when angler norms shift towards a total voluntary catch-and-release practice, even the best intended regulations may fail in producing trophy fish, particularly in top predators that show Ricker-type stock recruitment. Similarly, when stunting occurs below a minimum-size limit due to density-dependent growth, harvesting the stock to thin out individuals is recommended (FAO, 2012; Tesch, 1959), but this regulation often fails as anglers are not willing to keep very small fishes (Pierce & Tomcko, 1998). Clearly, appropriate fisher behavior is a necessary precondition that the harvest regulations achieve their intended objective. Our model did not explicitly model fisher behavior and thus the regulatory performance might not necessarily apply in real fisheries.

### Size-dependent maternal effects

Our model shows that the relative performance of minimum-length limits and harvest slots was largely robust to assumptions about size-dependent reproductive output, unless the mass-fecundity scaling was assumed extremely large at values only rarely reported in empirical studies of batch fecundity (Barneche et al., 2018). Importantly, however, we find that the well-established assumption of isometric scaling of mass and fecundity is already sufficient to justify increasing conservation of large fish through harvest slots when the numbers of fish harvested is to be maximized in addition to biomass yield. Our findings are thus in agreement with several previous studies revealing size-dependent maternal effects are not of sufficient importance to independently justify alternative size selectivity (Arlinghaus et al., 2010; Calduch-Verdiell, MacKenzie, Vaupel, & Andersen, 2014; McGillard, Punt, Hilborn, & Essington, 2017; O’Farrell & Botsford, 2006; Shelton et al., 2015). Framed differently, derivation of optimal size limit is largely robust against size-dependent maternal effects, particularly in relation to egg viability effects (that effects the slope of the stock-recruitment curve) and less so in relation to hyperallometry in fecundity (which strongly affects egg production and hence affects where on the x-axis one is on the stock-recruitment curve) (Figure 2). There was one notable exception - when hyperallometry in fecundity was strong, harvest slots became the optimal regulation at moderate to high fishing pressures when maximizing biomass yield under both Ricker and Beverton-Holt-stock recruitment, and the optimal harvest slot appeared earlier and narrowed when the goal was to maximize harvest numbers (Figure 8). Strong hyperallometry is most likely in batch-spawning species (Bernache et al., 2018; Marshall et al., in review). Specific for pike the maximum exponent of hyperallometry in fecundity reported so far is 1.22 (Arlinghaus et al. 2010) - a value where predictions of optimal size limits under isometry and hyperallometry did not differ much.

The latter result might sound surprising in light of several recent papers reporting that not accounting for hyperallometry in fecundity in classical fisheries models, when in fact it is present, will foster unsustainable overexploitation of very fecund, large individuals and strongly affect fisheries performance (Barneche et al., 2018; Cooper et al., 2013; Marshall, Gaines, Warner, Barneche, & Bode, 2019). The cited papers largely substantiate their conclusions based on comparisons of total egg output in populations fished with and without assumptions of hyperallometry in fecundity. Indeed, we also show that metrics that are sensitive to total egg production, e.g., spawning potential ratio (SPR), are strongly affected by hyperallometry in fecundity (Figure 2). This perspective, however, neglects the critical role of density dependent juvenile survival and growth compensation for ultimately affecting population dynamics and thus yield production or other fisheries outcomes. Our model does not focus on just egg production metrics as reference points and instead considers the entire life history, and the resulting productivity as a function of multiple sources of density-dependence. Importantly, we ask a different question - do assumptions of size-dependent maternal effect produce alteration of optimal size-based harvest polies in light of emerging population dynamical effects? The answer to this question is - not substantially, unless hyperallometry in fecundity is very high. We nevertheless recommend careful empirical estimation of mass-fecundity relationships if a model such as ours is to be used for concrete fisheries.

We found hyperallometry to start to matter for certain metrics (in particular yield and harvest numbers) when the scaling was very high (in our model 1.29 or higher). This is a very high value given the empirical evidence for batch fecundity. The average scaling of batch fecundity and mass in the study by Bernache et al. (2018) across a vast range of marine fish species is 1.18, and the maximum value across hundreds of marine fish species is 1.58. However, if repeat spawning is considered the mean value across 26 stocks rises to 1.89 (Marshall et al., unpublished data). Individual species such as *Engrualis mordax, Thunnus albacares* and *Sardinops sagaxcan* show scalings larger than 2 (Marshall et al., unpublished data). These values move the degree of hyperallometry into areas where harvest slots, rather than minimum-size limits, were predicted to maximize biomass yield. Given the existing evidence compiled by Marshall et al. (unpublished data), we expect hyperallometry to matter particularly in repeat spawners, to which pike do not belong. Specific for pike, Frost & Kipling (1967) reported isometric scaling of fecundity with mass. And although size-dependent maternal effects on egg viability were reported under controlled situations in a range of species (e.g., Berkely, et al. 2004b; Bravington et al., 2016; Venturelli et al., 2010), including pike (Arlinghaus et al., 2010), there is good reason to assume that such effects are an adaptation to specific natural environments. For example, an increased viability per egg in eggs spawned by large females may be an evolutionary adaptation to higher per capita fecundity in large individuals or an adaptation to significantly different spawning times (Marshall, Heppell, Munch, & Warner, 2010). Thus, it is unclear whether the optimistic scenario of size-dependent maternal effects modelled in our work would be widespread in nature. Independent of this uncertainty, the need to preserve large fish already emerges from isometric scaling of fecundity under certain objectives, e.g., when the target is to maximize harvest numbers or to achieve the best overall compromise regulation (see also Arlinghaus et al., 2010; Gwinn et al., 2015).

### Pretty good outcomes at the compromise harvest slot

We found that the optimal compromise regulation of a harvest slot achieved pretty good outcomes across all parameter combinations and management objectives, which were 50% or larger than the maximum possible single-objective outcomes at equilibrium. Under both Ricker and Beverton-and-Holt recruitment, in the compromise harvest regulation the biomass yield was the lowest of all possible outcomes. This means that the optimal harvest slot limit regulated the effective fishing mortality rate to levels lower than the fishing mortality rate at MSY. Earlier qualitative reasoning has argued that reducing fishing mortality rate below F_MSY_ would provide a “zone of new consensus” among traditionally conflicting conservation (erring to lower mortality rates) and fisheries objectives targeting MSY (erring towards more intensive harvesting rate, optimally Fmsy) (Hilborn, 2007). There is a long­standing debate that F_MSY_ should be considered the limit harvesting rate in fisheries rather than the target (Larkin, 1977) because of multiple risk of misspecifying F_MSY_ for single species in a community (Walters, Hilborn, & Christensen, 2008) as well as to avoid ecosystem and food web effects associated with highly size-truncated spawner populations (Francis, Hixon, Clarke, Murawski, & Ralston, 2007). Our model does not account for multi-species interactions, but multi-species models also suggest that the best compromise among fisheries and conservation can be achieved through fishing mortality rates smaller than F_MSY_ (Worm et al., 2009). A similar prediction is derived from size-spectrum models (Law & Plank, 2018). Our single species prediction of the compromise outcomes being a fishing mortality rate smaller than Fmsy thus agrees with alternative model formulations and perspectives that factor in other conservation targets (e.g., conservation of a more natural size and age structure) from an ecosystem or community-based harvesting perspective.

### Limitations

Our model has a number of limitations that could affect our results. Nine are worth mentioning.

First, we did not consider dynamic effort responses to the implementation of the harvest regulations and instead determined the optimal harvest regulation, given the objective and fixed maximum fishing mortality rates. The optimal harvest regulation effectively controlled fishing mortality. In real fisheries, fishers will respond to harvest regulations by altering behavior directly in response to the regulation (Beard, Cox, & Carpenter, 2003), might respond to changes in the fish stock and resulting expected catch rates or sizes or fish (Allen et al., 2013; Johnston et al., 2010) and possibly engage in non­compliance, particularly when catch rates drop (Johnston et al., 2015). These sources of implementation uncertainty can have far-reaching consequences for regulation performance in real fisheries (Allen et al., 2013, Johnston et al., 2015). They do, however, not fundamentally affect the conclusion of our equilibrium model as to the relative performance of different size-based harvest regulations given a certain objective.

A second limitation relates to the equilibrium nature of our analysis, which ignored temporal variation due to environmental stochasticity and transient dynamics. Several studies have shown that age truncation destabilizes stock dynamics in the face of environmental drivers unrelated to fishing (e.g., Anderson et al., 2008; Botsford et al., 2014; Hsieh et al., 2006; Ohlberger, Thackeray, Winfield, Maberly, & Vpllestad, 2014; Rouyer et al., 2011; Stige et al., 2017; Wikstrôm, Ripa, & Jonzén, 2011). One of the implied mechanisms relates to the fact that age truncation increases the non-linear dynamics of population growth and hence subsequent years of poor recruitment due to environmental forcing can have strong “resonance” in population dynamics and destabilize abundances (Botsford et al., 2014). If these processes occur, it would reinforce the key findings of the present model as harvest slots outperformed minimum-length limits as a compromise regulation, which reduce juvenescence effects and maintain old age structure (see also LeBris et al., 2014). However, our analysis should depict long-term average responses of populations to each management strategy and be robust to such a long-term perspective.

Third, although we carefully accounted for fecundity and viability benefits of large spawner size, the model omitted other reproductive processes related to body size, e.g., different spawning times by differently sized fishes, the ability of large fish to lead spawning migrations or otherwise affect reproductive output, e.g., through size-based sexual selection (Hixon et al., 2014; Jprgensen, Dunlop, Opdal, & Fiksen, 2008; Uusi-Heikkilâ, Bôckenhoff, Wolter, & Arlinghaus, 2012). Our model is thus conservative as to the conservation benefits of large spawner size.

Fourth, although we carefully included key aspects of density and size-dependency, it is possible that density effects also occur on fecundity itself (as for example shown in pike, Craig & Kipling, 1983). An age-structured model including this process in northern pike, essentially reported similar findings to the present study (Arlinghaus et al., 2010), so we conclude the omission of density-dependence in fecundity is unlikely to fundamentally alter the present conclusions.

Fifth, we present a single-species model, and naturally a target species might be affected by multiple ecological processes beyond the single species demography. However, other work using community based size-spectrum models, which represent community dynamics, similarly suggest that harvest slots may lead to more balanced fishing than classical “knife-edge” selectivity through minimum-length limits (Law & Plank, 2018).

Sixth, our model assumed full compensation among age classes in terms of density-dependent growth, i.e., all individuals of all ages relied on the same prey resources. While this assumption may hold for pike who become piscivorous in the first year of life (Persson et al., 2006) and feed on similarly sized prey fish as they age (Gaeta et al., 2018), other species show more complex ontogeny in prey choice and thus the competition for food will vary strongly by size(age) class (e.g., in Eurasian perch, *Perca fluviatilis,* Claessen et al., 2004; Persson et al., 2003). Size-structured models are needed to account for more complex food-dependent growth and size-dependent interactions where different cohorts feed on different prey types (van Kooten, Persson, & de Roos, 2007).

Seventh, we also assumed all individuals to remain fully vulnerable even after being released. There is increasing understanding that fish learn to avoid being recaptured after initial private hooking and release experiences (e.g., Klefoth, Pieterek, & Arlinghaus 2013). Also, gear avoidance behavior is reported for a range of commercial fishing gears (Arlinghaus et al., 2017). It is particularly the largest and oldest individuals that reduce their vulnerability to the gear over time. Such behavior would naturally create a “harvest slot” or dome-shaped selectivity as reported for angling gear in other studies (O’Farrell & Botsford, 2006).

Eighth, we assumed fish to maintain reproductive performance even at old ages, thereby assuming no reproductive senescense. Reproductive senescence has been reported in viviparous (Reznick, Bryant, & Holmes 2004) as well as broadcast spawners (Benoit et al., 2018) and has been implicated in causing reduced recruitment in unexploited esocid stocks (Eslinger, Dolan, & Newman, 2010; Eslinger, Sass, Shaw, & Newman, 2017). However, reproductive senescence is unlikely to affect substantial numbers of fish in exploited stocks and therefore is unlikely to matter much for popluations dynamics. Indeed, a model by Arlinghaus et al. (2010) specific for pike assumed the presence or absence of reproductive senecense revealing negligible impacts of reproductive senescence on the performance of size limits.

Finally, we did not consider the potential for fisheries-induced evolution (FIE). Although FIE can shift maturation size and age, elevate reproductive output and reduce post maturation growth (Jprgensen et al., 2007), several models have shown that the relative phenotypic change expected within the realm of plasticity is orders of magnitude greater than life-history change caused by selection (Eikeset et al., 2016; Lester et al., 2014). Studies specifically focusing on addressing FIE have also shown that keeping fishing mortality rates within limits that optimize ecological targets (e.g., MSY) are usually also sufficient to address FIE (Eikeset, Richter, Dunlop, Dieckmann, & Stenseth, 2013). Previous work has also shown that harvest slots can ameliorate key selection responses in life-history traits (Matsumura et al., 2011; Zimmermann & Jørgensen, 2017), reinforcing our study findings.

### Conclusions and implications

Our work suggests that harvest slots were often the superior regulation to the classical minimum-length limits across the suite of yield and catch-based management objectives considered, particularly when representing density-dependent growth, size-dependent mortality and hyperallometry in fecundity and more fisheries objectives than biomass yield alone. Our results suggest that even for biomass yield where minimum-length limits are typically considered optimal, we found harvest slots to produce yields similar to values predicted from the optimal minimum-size limit. Harvest slots also turned out to consistently constitute the best regulation when considering the full suite of catch and yield based objectives. Thus, depending on the management objective harvest slots can be recommended as a suitable alternative to minimum-length limit, in agreement with several previous studies (Ayllón et al., 2019; Arlinghaus et al., 2010; García-Asorey, Escati-Penaloza, Parma, & Pascual, 2011; Gwinn et al., 2015; Jensen 1981; Koehn & Todd, 2012; Le Bris et al., 2015, 2018; Reed, 1980). Because we found our results to be fairly insensitive to most input parameters and the different compensatory processes we modelled (e.g., stock-recruitment function), we are confident that our results hold for a wide range of species. Clearly, if very large fish have high value as a landed (as opposed to just captured) fish, the predictions of our model would not hold as one of the key objectives we optimized was the catch, not the harvest, of particularly large (trophy) fish. Harvest slots may also reduce the recovery speed as the exploited stock is composed of slower growing (large) individuals relative to stocks expected under minimum-length limit regulations (LeBris, Pershing, Hernandez, Mills, & Sherwood, 2015). Ultimately, our results suggests the choice of the local size limits will crucially depend on management objectives, local fishing pressure, the stock-recruitment function and the density and size-dependency of growth, mortality and fecundity.

From a practical perspective, compromises among multiple yield and catch-based objectives that favor harvest slots over minimum-length limits are likely to be made in most mixed commercial-recreational fisheries, e.g., for the mixed fishery in northern pike in the Baltic, as well as for the prototypical recreational fishery where not only harvest but also the catch of large fish or catch rates are part of the utility function (Arlinghaus, Beardmore, Riepe, Meyerhoff, & Pagel, 2014; Beardmore et al., 2015). Harvest slots could be straightforwardly implemented in small-scale fisheries where gear types such as fyke nets, gill nets (with regulations on lower and upper mesh sizes) or long-lines are prevalent (assuming captured fish can be released unharmed) and the catch of immature and very large individuals can be either controlled or protected sizes be released unharmed. Harvest slots are particularly easily implemented in recreational fisheries (Gwinn et al., 2015; Tiainen et al., 2017). In order to function, the discard mortality has to be controlled and the width of the harvest slot tuned to local fishing pressures.

## Acknowledgements

We thank Dan Gwinn and Kai Lorenzen for discussions in an early phase of this project and Philipp Czapla for help with formatting. We thank Ian Winfield, Yngvild Videnes and 0ystein Langangen for support with unpublished data used to derive parameters included in this model. Funding was received by RA through the European Maritime Fisheries Fund (EMFF) of the EU and the State of Mecklenburg-Vorpommern (Germany) (grant MV-I.18-LM-004, B 730117000069). Further funding was received by the University of Florida to support a sabbatical of RA in Gainesville in 2013 where this project started and by the Leibniz-lnstitute of Freshwater Ecology and Inland Fisheries supporting a guest scientist stay by RNA in Berlin in 2019, during which this manuscript was completed. There is no conflict of interest.

